# CRISPR/Cas9-Based Edition of Frataxin Gene in *Dictyostelium discoideum* for Friedreich’s Ataxia Disease Modeling

**DOI:** 10.1101/2023.02.27.530330

**Authors:** Hernan G. Gentili, María Florencia Pignataro, Justo Olmos, Florencia Pavan, Itati Ibáñez, Javier Santos, Duarte Francisco Velázquez

**Affiliations:** Instituto de Biociencias, Biotecnología y Biología Traslacional (iB3). Departamento de Fisiología y Biología Molecular y Celular, Facultad de Ciencias Exactas y Naturales, Universidad de Buenos Aires. Intendente Güiraldes 2160, Ciudad Universitaria, C1428EGA, Buenos Aires, Argentina; Instituto de Química Física de los Materiales, Medio Ambiente y Energía (INQUI- MAE), CONICET, FCEN, UBA, Intendente Güiraldes 2160, Ciudad Universitaria, C1428EGA, Buenos Aires, Argentina; Departamento de Química Biológica, Facultad de Ciencias Exactas y Naturales, Universidad de Buenos Aires. Intendente Güiraldes 2160, Ciudad Universitaria, C1428EGA, Buenos Aires, Argentina

**Keywords:** Iron-Sulfur Cluster Assembly/ Frataxin/ Friedreich Ataxia/Rare Disease

## Abstract

In this paper we describe the development of a new model system for Friedreich’s Atax- ia (FA) using *Dictyostelium discoideum*. We investigated the conservation of function between humans and *D. discoideum* and showed that DdFXN can substitute the human version in the interaction and activation of the Fe-S assembly supercomplex. We edited the *fxn* locus and isolated a defective mutant, clone 8, which presents landmarks of frataxin deficiency, such as a decrease in Fe-S cluster-dependent enzymatic functions, growth rate reduction, and increased sensitivity to oxidative stress. In addition multicellular development is affected as well as grow on bacterial lawn.

We also assessed the rescuing capacity of DdFXN-G122V, a version that mimics a human variant present in some FA patients. While the expression of DdFXN-G122V rescues growth and enzymatic activity defects, as DdFXN does, multicellular development defects were only partially rescued

The results of the study suggest that this new model system offers a wide range of pos- sibilities to easily explore diverse phenotypes in FA and develop drug or treatment screenings for designing and evaluating therapeutic strategies.

## Introduction

*Dictyostelium discoideum* is a social amoeba that lives in the soil and feeds on bacteria and other microbes. Dictyostelids belong to a separate branch of eukaryotic organisms, distinct from plants, fungi and animals. Its cells lack a cell wall resembling animal cells in organization (*1*). *D. discoideum* has become a very attractive eukaryotic non-mammalian model organism to study molecular mechanisms, cell physiology and human pathology (*2, 3*). The studies carried out with *D. discoideum* have provided great insights into diverse areas such as bacterial infection, immune cell chemotaxis, and autophagy/phagocytosis as well as mitochondrial and neurological disorders (*4*).

This amoeba has been widely used to study human diseases, including neurodegenerative illnesses such as Alzheimer’s, Parkinson’s, Huntington’s, as a result of which important discoveries of the pathophysiology of these pathologies have been obtained (*5*). Furthermore, *D. discoideum* has been used as a model to identify drug targets and discover new compounds with therapeutic potential, and these advances have even served as a platform for new clinical trials (*6, 7*). This organism allows studying the effect of very specific molecular alterations, from different viewpoints, combining biochemistry, structural biology, and cell biology to generate a comprehensive picture concerning the outcomes of the alteration at different levels of complexity. *D. discoideum* has a unique developmental life cycle among eukaryotes, presenting both unicellular and multicellular phases. This allows the study of different levels of cell organization and cell-cell interaction and communication (**Figure 1**).

**Figure 1.**
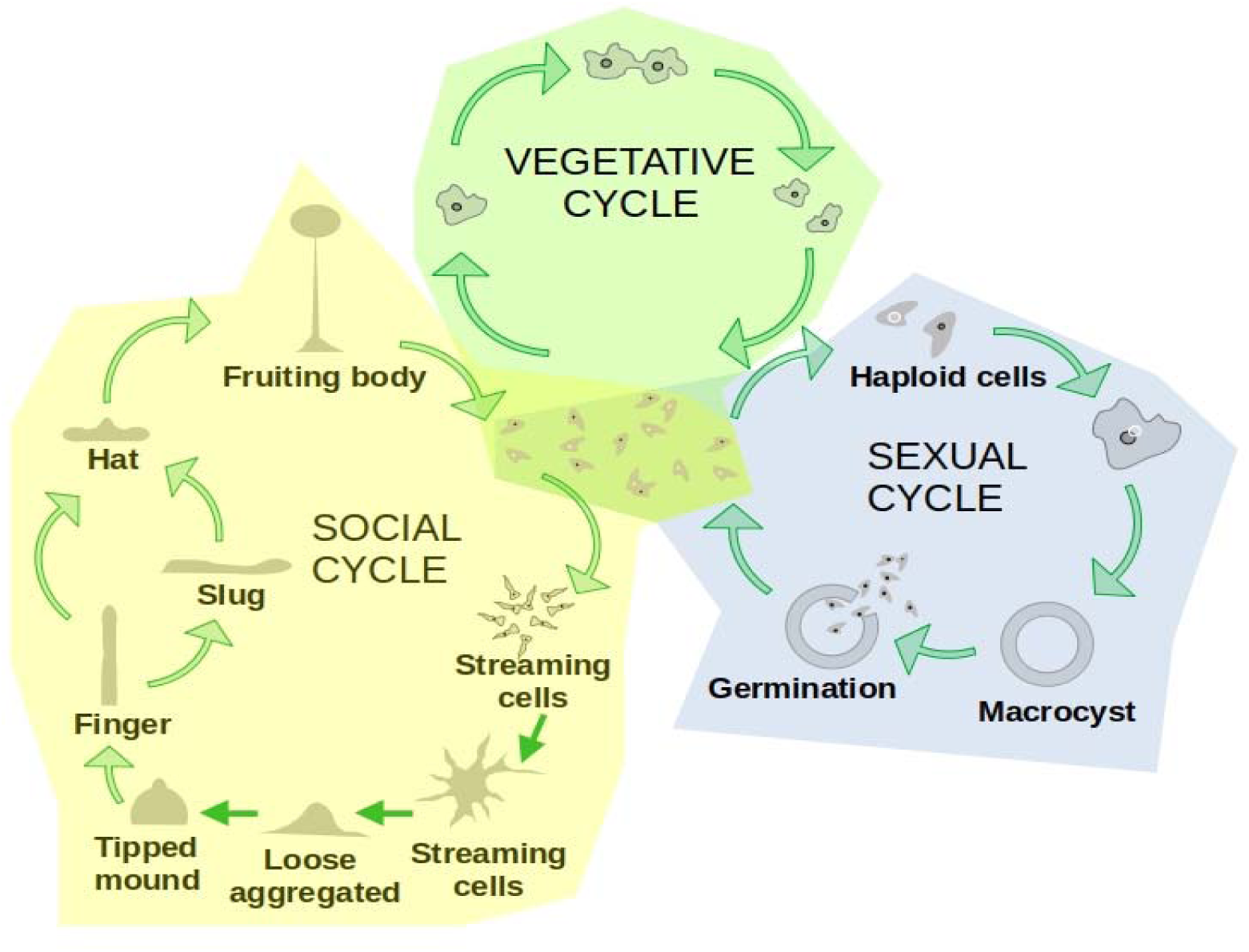
*D. discoideum* Life Cycle. *D. discoideum* grows as unicellular amoebae as long as there are nutrients available (Vegetative Cycle). During this growth phase, haploid amoeboid cells feed on bacteria and divide by a binary fission process. In response to starvation, *D. discoideum* triggers a multicellular developmental program that leads to the formation of a fruiting body bearing a mass of spores for dispersal (Social Cycle). During this program, starved cells gather by chemotaxis and form an aggregate. This aggregate suffers several morphogenetic changes going through different stages. First, cells form a mound, enclosed within an extracellular matrix. After that, the mound develops into a standing structure (finger). The finger may develop into a migrating slug, or it may directly progress all the way to the culmination stages. Finally, the fruiting body is formed. It will contain spores that will be released and eventually germinate to produce growing amoeboid cells (*8*). Alternatively, upon starvation, *D. discoideum* can go through a sexual cycle if sexually compatible cells are present and other environmental conditions are met (*9*).

Recently, CRISPR/Cas9-mediated technology has been implemented to *D. discoideum*, allowing rapid genome editing by transiently expressing single guide RNA (sgRNA) and Cas9 using an all-in-one vector and the generation of genomic mutants (*10–12*). This possibility may accelerate gaining knowledge of specific components of the molecular machinery underlying the complex life cycle in *D. discoideum*.

In addition to what was stated above, *D. discoideum* helps to implement the “Three Rs principle” in animal research (Replacement, Reduction, and Refinement); so development of *Dictyostelium*–based disease models will be highly beneficial in preliminary drug screenings (*13*). D. *discoideum* has been identified as an exceptional model organism to study rare diseases (*14, 15*). More than 8000 different pathologies, affecting more than 8% of the world human population, have been classified to date as rare diseases; they are generally underrepresented in scientific agendas and suffer from a lack of tools, model organisms and therapeutics.

In this context, our laboratory is interested in the development of new tools to facilitate the search for therapeutic solutions for Friedreich’s Ataxia (FA), a neurodegenerative disease that affects 1:50000 of the population worldwide. The main cause of this disease is a decrease in the expression or in the functionality of a mitochondrial protein named frataxin, which is encoded in the nuclear genome.

Frataxin is involved in the biosynthesis of the iron-sulfur cluster (Fe-S) in the mitochondrial matrix. The deficiency of frataxin affects several enzymatic reactions that depend on Fe-S clusters, which are essential cofactors involved in several enzymatic functions. The Krebs cycle enzymes, aconitase and succinate dehydrogenase (*16*), and the respiratory chain complexes (*17, 18*); critical processes such as DNA repair (*19*) or chemical modification of transfer RNAs (*20*); lipoic acid synthase, which catalyzes the final step in the novo pathway for the biosynthesis of lipoic acid (a key coenzyme of pyruvate dehydrogenase and the α-ketoglutarate dehydrogenase enzymes) (*21*), all depend on Fe-S clusters.

The Fe-S assembly reaction depends on a supercomplex formed by at least five different subunits: i) the L-Cys desulfurase NFS1, a pyridoxal-phosphate (PLP)– dependent enzyme, which catalyzes the desulfurization of L-cysteine, generating as products the precursor sulfide attached as a persulfide group to a Cys residue (Cys-S-SH) and L-alanine; ii) the ISD11 protein, which is only present in eukaryotic organisms (*16, 22*); iii) the mitochondrial acyl carrier protein (ACP), which stabilizes ISD11(*23, 24*); iv) the scaffolding protein (ISCU); and v) frataxin.

The NFS1 dimer is stabilized by the ACP-ISD11 heterodimer (*25*). The Fe-S cluster assembly site is situated on the scaffolding protein ISCU and not only is frataxin the kinetic activator of the reaction, but also its surface is part of the Fe-S cluster assembly. The stoichiometry of the supercomplex is (NFS1-ACP-ISD11-FXN-ISCU)_2._ Its architecture is intricate as each heterodimer ACP-ISD11 also interacts with both NFS1 subunits as a bridge and two assembly sites formed by ISCU/FXN and both NFS1 chains. That is, frataxin simultaneously interacts with both NFS1 subunits and ISCU (*26*). Even though it has been previously demonstrated that frataxin binds iron, it is not clear whether it works as a chaperon of this metal ion in the context of the supercomplex. It has also been shown that ISCU is able to bind this metal and that this activity is crucial for Fe-S cluster assembly (*27*).

In addition to having Friedreich’s Ataxia caused by mutations in frataxin gene, the alteration of transcription, aberrant splicing or the presence of point mutations in other proteins involved in the cluster assembly and affecting their expression or functionality all result in severe human diseases. Thus, the mutation of NFS1 results in an autosomal recessive mitochondrial disease characterized by a respiratory chain complex II and III deficiency and multisystem organ failure (*28*), and the mutation of ISCU results in ISCU myopathy (*29*), whereas the mutation of ISD11 (R68L) is associated with the development of a mitochondrial genetic disorder, i.e., an autosomal recessive disease, known as Combined Oxidative Phosphorylation Deficiency 19 (COXPD19) (*17*).

The mutations of frataxin that result in FA affect the protein at different levels: conformational stability (e.g., L198R, G137V, G130V) (*30–32*), the mitochondrial import and processing pathway (e.g., W168R and W173G) (*33*), iron binding affinity (e.g., D122Y) (*32, 34*) or alterations at the assembly site architecture (e.g., W155R and N146K and Q148R) (*35, 36*). In addition, the truncation of frataxin at position 193 (deletion of the last stretch of residues named the C-terminal region, which conforms a non-periodic structure) results in a pathogenic variant (*37*). A very similar variant, truncated at position 195, exhibits strong alterations in its internal motions and also reduced conformational stability, besides a decrease in its iron binding capability (*30*).

Frataxin has been described as an essential protein in eukaryotic organisms as the deletion of this protein is lethal in yeast and mammalian cells (*38*). Moreover, in multicellular eukaryotic organisms (plants and mice), the complete deletion of frataxin leads to early embryonic lethality (*39, 40*) or to the arrest of the larval stage L2 / L3 in *Caenorhabditis elegans* (*41*) and reduced larval viability along with metamorphosis failure in *Drosophila melanogaster* (*42, 43*). Regarding the use of cell cultures, different models have been described. Since the patient’s cell line (fibroblasts or lymphocytes) does not consistently express the biochemical phenotypes associated with FA under basal culture conditions, RNA interference (RNAi) strategies have been developed to reproduce partial frataxin deficiency in human and murine cell lines (*44, 45*). Furthermore, “humanized” murine cell models have been developed to eliminate endogenous frataxin and express frataxin with pathogenic mutations (*46*). A murine fibroblast cellular model, with the ability to deactivate frataxin transcription, was also generated using the Cre /loxP recombination system. On the other hand, stem cells from patients have been used, whereas the use of inducible pluripotent stem cells that can mimic tissues affected by FA is under development (*47, 48*).

In a previous paper we examined the *D. discoideum* genome and found the complete dotation of proteins involved in the Fe-S cluster assembly (*49*). Furthermore, by analyzing the sequences and structure models of these proteins, we found that residues located in the protein-protein interaction surfaces are highly conserved between the amoeba and the human. In fact, the frataxin residues involved in FA are fully conserved, with the exception of a core residue His183 of the human FXN that interacts with residues located in the CTR (the last stretch of the protein), whereas in DdFXN, this residue is an Arg fully exposed to the solvent, and a Tyr residue (in DdFXN) is located at the position corresponding to the Trp168 in the human variant, as inferred by means of a structure model of DdFXN.

In this paper, we investigated the effect of the functional deficiency in frataxin on the metabolism and physiology of the amoeba *D. discoideum*. The short duplication time, simple genetic manipulation, the simplicity of creating knockout cell lines by CRISPR/Cas9 due to its haploid genome and the simplicity of the characterization of phenotypes were conceived as key advantages of this model.

Some of the alterations observed in mammalian cells are also present in *D. discoideum*, such as reduced aconitase and succinate dehydrogenase or a higher sensitivity to oxidative stress. In addition, the amoeba cultures deficient in frataxin grew at a lower rate and the life cycle exhibited alterations. In addition, we have explored the rescue capacity of the constitutive frataxin expression.

These cell lines obtained by CRISPR/Cas9 will allow us to carry out therapeutic screenings of different compounds and drugs that can be used in the treatment of FA.

## Material and Methods

### Strains, Cell Culture, Plasmid Constructions, CRISPR/Cas9 Guide Design and D. discoideum Transformation

The *D. discoideum* strain AX2 was cultured at 22 °C on axenic culture on dishes, in bacterial medium SM agar plates with *Klebsiella aerogenes* lawn, or in Erlenmeyer flasks, in HL5 medium (http://dictybase.org/techniques/index.html). The parental plasmid used for CRISPR/Cas9 genome editing (pTM1285) (*50*) was kindly provided by Dr. Kamimura. It was used to insert the DNA fragment codifying for the RNA guide to target the *D. discoideum fxn* gene. For the guide preparation, a pair of oligonucleotides (FwGuide553/RevGuide553) designed using CRISPOR (http://crispor.tefor.net) and Breaking-Cas (https://bioinfogp.cnb.csic.es/tools/breakingcas/index.php) were adequately annealed, and for DNA ligation, the Golden Gate strategy was used (**Table 1**). Construct checking was performed by colony PCR of *E. coli* DH5α, using the RevGuide553 and tRNA_seq_3 as primers.

**Table 1.**
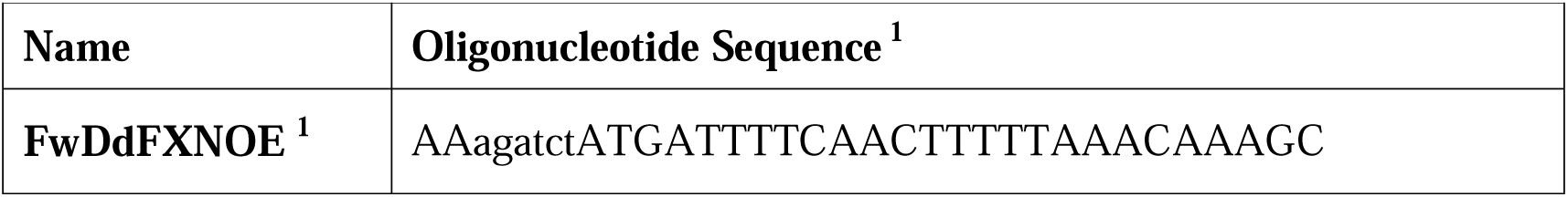
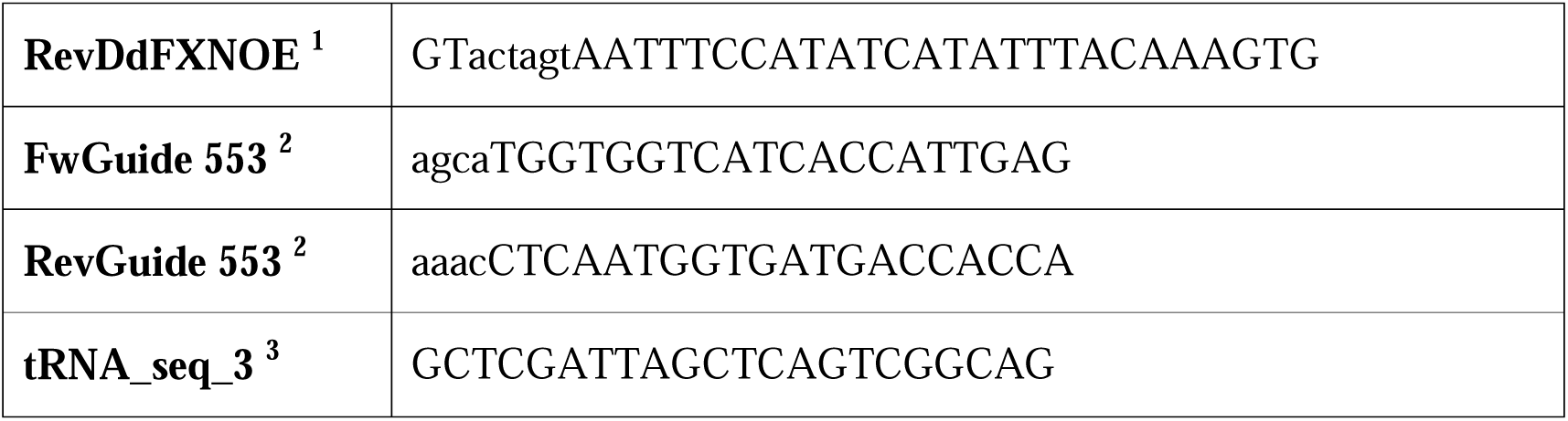
Oligonucleotides Used in this Research. ^1^ Upper-case letter indicates match and lower-case mismatch 307 with target template.

For the transformation of *D. discoideum*, AX2 cells were harvested during the exponential phase of growth, washed twice in ice-cold H50 buffer (20 mM HEPES, pH 7.0, 50 mM KCl, 10.0 mM NaCl, 1.0 mM MgSO_4_, 5.0 mM NaHCO_3_ and 1.0 mM NaH_2_PO_4_), and re-suspended in H50 buffer at a concentration of 5×10^7^ cells/ml. A volume of 100 µl of cell suspension was electroporated using 10-20 µg of plasmid (0.75 kV, 25 µF, twice). Cells were transferred to culture dishes containing HL5 culture medium and incubated for 16 h. For clone selection, G418 antibiotic (geneticin), which blocks polypeptide synthesis, was added (20 µl/mL) and the cells were incubated for 48h; the resistance to G418 is conferred by the neo gene located in the pTM1285 vector.

After that, HL5 medium containing G418 was removed from the plastic dishes and the cells were re-suspended in a volume of 400 µL of HL5 medium; a volume of 75 µL was mixed with 250 µL of bacteria *Klebsiella aerogenes* liquid culture and plated in SM agar plates. After an incubation of 3-4 days at 22 °C, plaques were observed (absence of bacterial lawn).

### D. discoideum Clone Isolation and DNA Sequencing

To identify sgRNA/Cas9 editing, the isolated genomic DNA was directly carried out by picking up cell material from the corresponding *D. discoideum* plaque using a sterilized tip. The material was re-suspended in a volume of 50 µL of Lysis Buffer (50 mM KCl, 10 mM Tris-HCl pH 8.3, 2.5 mM MgCl_2_, 0.45% NP40, 0.45% Tween-20 and Proteinase K (40 µg). The cell suspension was incubated at 20-24 °C for 20 min and then heated at 95 °C for 3 min to inactivate the protease. The cell lysate was used as template for PCR. The DNA corresponding to the *fxn* gene was amplified using KOD hot start master mix (Cat. N° 71842 Millipore) and FwDdFXNOE and RevDdFXNOE as primers (**Table 1**). The PCR cycle comprised denaturation at 95 °C for 30 s, annealing at 55 °C for 30 s, an extension at 68 °C for 30 s, and then 40 PCR cycles were programed. The amplified DNA fragments were purified using Wizard SV Gel and the PCR Clean-Up System (Cat. N° A9281 Promega), then the DNA was sequenced using the Macrogen facility (https://dna.macrogen.com/).

### Evaluations of the D. discoideum Growth Rate and Life Cycle Alterations

To evaluate the generation time (*g*), *D. discoideum* growth curves (22 °C and 180 rpm) were made using flasks (250 mL) containing HL5 culture medium (75 mL) and starting from a 1×10^5^ amoeba cells/mL inoculum. The cell cultures were grown during a week, and small aliquots (0.1 mL) were regularly taken for cell counting using a Neubauer chamber. The growth curves were analyzed, and the maximum (μ) was obtained from the slope when the culture grows exponentially. The generation time was calculated according to Equation 1.

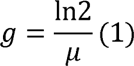

For the study of the *D. discoideum* life cycle and development, a 6-cm petri dish was prepared with 10 mL of 1.8% Oxoid L28 agar in KK2 medium (16.5 mM KH_2_PO_4_, 3.9 mM K_2_HPO_4_, pH: 6.2) and 2.0 mM MgSO_4_, 0.1 mM CaCl_2_ (complete KK2).

A volume of 5-15 mL of cell culture in the log phase (2-5 × 10^6^ cells/mL) was centrifuged at 1400 rpm for 3 min under and then two washes with KK2 were carried out. Next, the cells were resuspended in complete KK2 to give 2.5 × 10^7^ cell/mL. After that, a volume of 1.6 mL containing 4 × 10^7^ cells was added to each plate, and the plates were left in a levelled table for 15 min for cell adhesion, after which the medium was carefully removed by tilting the plates. The plates were incubated in humidity at 22 °C for 18-22 h.

### Aconitase and Succinate Dehydrogenase Activities Measurements

Aconitase (ACO) and succinate dehydrogenase (SDH) enzymatic activity measurements were carried out on the soluble fraction of the cellular extracts of *D. discoideum* AX2 and **clone 8**.

For the SDH activity assay, 2 × 10^5^ cells were used per reaction (in multi well plates) and cell lysis was performed by three freeze-thaw cycles. The assay was carried out according to the supplier’s instructions (Abcam: ab 228560).

To measure ACO activity, 1.5 × 10^6^ cells were used per reaction. In this case, cell lysis was performed by means of a detergent solution included in the commercial kit. The protocol was according to the supplier’s instructions (Abcam: ab 109712).

### Frataxin Variants Molecular Dynamics Simulation

Molecular dynamics simulations and the analysis of production runs were carried out using the YASARA Structure (*51*) on the following hardware: Processor Intel CORE i7 10,700 10th generation; SOCKET 1200 2.9 GHZ (Max 4.8 GHZ) 16 M cores/threads 8/16, 2; Memory Kingston HX426C16FB3/8G HyperX2666 MHz; Disc SSD Kingston A400 240GB SATA 7 mm; Linux Ubuntu 20.04 LTS 64 bit. The molecular models corresponding to wild-type DdFXN and a DdFXN fragment from clone 8 strain were modeled using AlphaFold2 (*52*) and with SwissModel at https://swissmodel.expasy.org/interactive.

The coordinates were solvated, and standard minimization protocols were applied to remove steric clashes. The simulation cell was prepared by maintaining a 20 Å water-filled space around the protein with a density of 0.997 g/mL. The system (cubic cell, periodic boundaries, and an 8.0 Å cut-off for long-range coulomb electrostatics forces) was neutralized with 0.9% NaCl, and the temperature was maintained at 298 K with a pH of 7.4. After the initial steepest descent minimization, unrestrained replicas of 100 ns MD simulations using an ff14SB Amber force field were carried out with 2.50 fs time steps (*53*). Snapshots were saved every 0.1 ns. The root-mean square deviation (RMSD), root mean square fluctuation (RMSF), and secondary structure content were calculated. Molecular dynamics simulations were performed using the Yasara Structure program (*54*).

### Western Blotting Analysis

For frataxin detection in cell lysates of *D. discoideum*, we used a set of nanobodies prepared in our laboratory using recombinant DdFXN as the target and phage display technology. Three panning rounds with increasing washing steps (10, 15 and 25 for rounds 1, 2 and 3, respectively) were carried out to select specific nanobodies. Since the nanobodies carry a 6xHis tag, we used anti RGS HIS6 HRP(QIAGEN) Cat.n°/ID 34450.

For the detection of DdFXN, wild-type and the G122V variant, an anti-FLAG monoclonal antibody (Cell Signaling Technology, [9A3] 8146S) was used followed by a secondary HRP-conjugated anti-mouse (ThermoFisher Cat. #31430) to specifically detect frataxin expressed from plasmidic DNA. For this detection, the FLAG of sequence DYKDDDDK was included in the C-terminal stretch of DdFXN. The HRP signal was detected using Clarity^TM^ Western ECL subtrate (Bio-Rad #1705060).

### Oxidative Stress Sensitivity Assay

To assess sensitivity to oxidative stress, cell viability was determined upon H_2_O_2_ treatment using a crystal violet assay described by Feoktistova et al. (*55*). In summary, 100000 cells were seeded in a 24-well plate and allowed to adhere for 1h. Then the medium was removed and fresh HL5 medium, or supplemented with 2 mM H_2_O_2_, was added to each well. Since adherent cells detach from cell culture plates during cell death, the wells were gently washed to remove the dying cells after the corresponding time of treatment. Remaining cells were fixed and stained with crystal violet. Culture plates were dried, and crystal violet was measured after solubilization with acetic acid (A570 nm) as an estimate of the remaining cells. Viability was calculated referenced to the crystal violet at time 0 for each strain. The experiment was performed at least 3 times for each strain and contained 4 replicates of each time point.

## Results

### DdFXN Can Substitute Human Orthologue in the Supercomplex for Fe-S cluster Assembly

Since we aimed to establish *D. discoideum* as a model organism for FA using DdFXN to explore its roles within the cell and to extrapolate it to the human context, we first tested the ability of DdFXN to substitute the human orthologue in the context of the supercomplex.

As proof of functional conservation, we assessed the ability of DdFXN to interact with the human supercomplex NIAU (NFS1/ACP-ISD11/ISCU)_2_. We carried out fluorescence anisotropy assays using a variant of DdFXN (DdFXN_C35A) or human FXN S202C labelled with Texas-red.

The labelled frataxin variant, in the presence of saturating ISCU concentration, was incubated with increasing NIA concentrations. As it can be seen in **Figure 2A**, the DdFXN variant was able to bind to the human NIAU in a similar fashion as the mature form of the human FXN does, although with a lower apparent affinity. It is worthy of note that the wild-type DdFXN was able to compete with the labeled-human-frataxin variant (**Figure 2A, inset**), suggesting that the same binding site is involved for both proteins.

**Figure 2.**
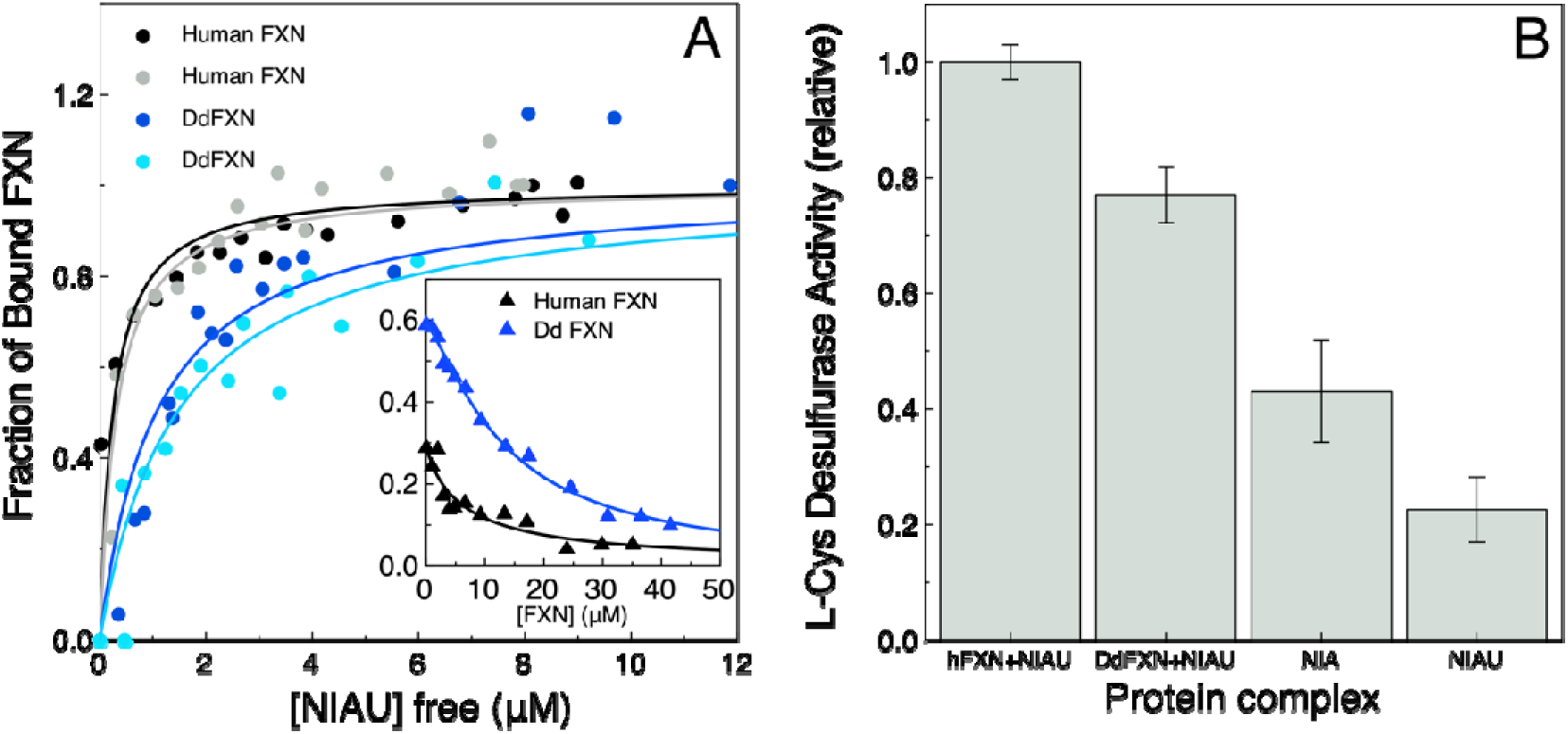
In Vitro Substitution of Human FXN by DdFXN. (A) DdFXN interaction with the human supercomplex L-Cys desulfurase. Texas red-Labeled DdFXN (blue and cyan correspond to two independent experiments) or human FXN (black and gray correspond to two independent experiments) were incubated with increasing concentrations of the NIAU subcomplex and fluorescence anisotropy was monitored. A fraction of bound frataxin was plotted as a function of the free NIAU complex. The inset shows the competition with non-labeled proteins of a preformed supercomplex in which human FXN was labeled with Texas red followed by fluorescence anisotropy. The competition was carried out using non-labeled human FXN (black triangles) or non-labeled DdFXN (blue triangles); the fraction of bound frataxin was plotted as a function of the added FXN variant. (B) In vitro activity of the human supercomplex activated by human FXN or DdFXN. The activity corresponding to NIA or the NIAU complexes without frataxin are shown for reference.

We then tested whether this binding is functional and if DdFXN is also able to act as the kinetic activator of L-Cys-desulfurase activity in a human supercomplex context. **Figure 2B** shows that the interaction of DdFXN with the human supercomplex was productive and Cys-desulfurase activity increased upon the addition of human FXN or DdFXN.

### Fxn Locus Can Be Easily Edited in D. discoideum with CRISPR/Cas9 Technology

As noted above, we aimed to generate a new experimental model to study the biochemical consequences of alterations in the functionality of frataxin in the cell. We intended to get mutant strains that were highly deficient in frataxin so we would be able to explore strategies for reestablishing homeostasis and to evaluate the rescue capacity of wild-type and disease-associate variants. With this aim in mind, we carried out CRISPR/Cas9 editing of *D. discoideum* endogenous locus, using a guide RNA targeted at Trp146. This residue, corresponding to the human Trp155, is extremely conserved in frataxin along the tree of life because of a functional role (*49*). When looking at the structure of the human supercomplex (NFS1/ACP-ISD11/ISCU/FXN)_2,_ Trp155 is located near the docking surface of ISCU and at Van der Waals distance of its [2Fe-2S] assembly site, and from Leu386 of NFS1, which hints at its importance. Trp155 belongs to a highly conserved region that contains the Motif 1 described by Gibson *et al*. (*56*).

Hence, it would be more likely to obtain deleterious mutations in this region. Using pTM1285, an all-in-one vector, to express the corresponding guide RNA, we obtained hundreds of clones and selected 14 presumably edited clones (**Figure S1**).

After the isolation of these clones, the genomic DNA corresponding to the *fxn* gene was successfully amplified and sequenced for 13 clones. The analysis of the genomic DNA sequences showed a variety of mutations in the protein as the result of CRISPR/Cas9 editing and subsequent DNA repair (**Table 2**, **Figure S1**).

**Table 2.** Summary of CRISPR/Cas9 Edition of fxn Locus in D. discoideum in Sequenced Clones. ^1^ Seven different editions.

| Mutation | Number of Clones |
| --- | --- |
| Premature STOP codon | 9 (7)<sup>1</sup> |
| Small Insertion | 2 |
| Point Mutation | 1 |
| Silence Mutation | 1 |

<sup>1</sup> Seven different editions.

### A D. discoideum Clone Lacking Frataxin is Viable

The amino acid sequence analysis of *D. discoideum* clones obtained by CRISPR/Cas9 genome editing (*e.g.*, **clone 8**) indicated that *fxn* gene functionality of DdFXN can be obliterated without being lethal for the amoeba, at least in the experimental conditions assayed in this paper.

Edition in **clone 8** produces a large truncation of the protein chain because of one base pair deletion generating a frame shift, and premature stop codon (**Figure S2**). This truncation eliminates a stretch that includes more than 40% of the sequence, which may completely destabilize the structure of the remaining protein fragment, comprising residues 81-150 from DdFXN, using the numbering of the DdFXN precursor (**Figures 3A and B**). The truncation eliminated part of strand beta 4, strands beta 5 and 6, alpha helix 2 and the CTR.

**Figure 3.**
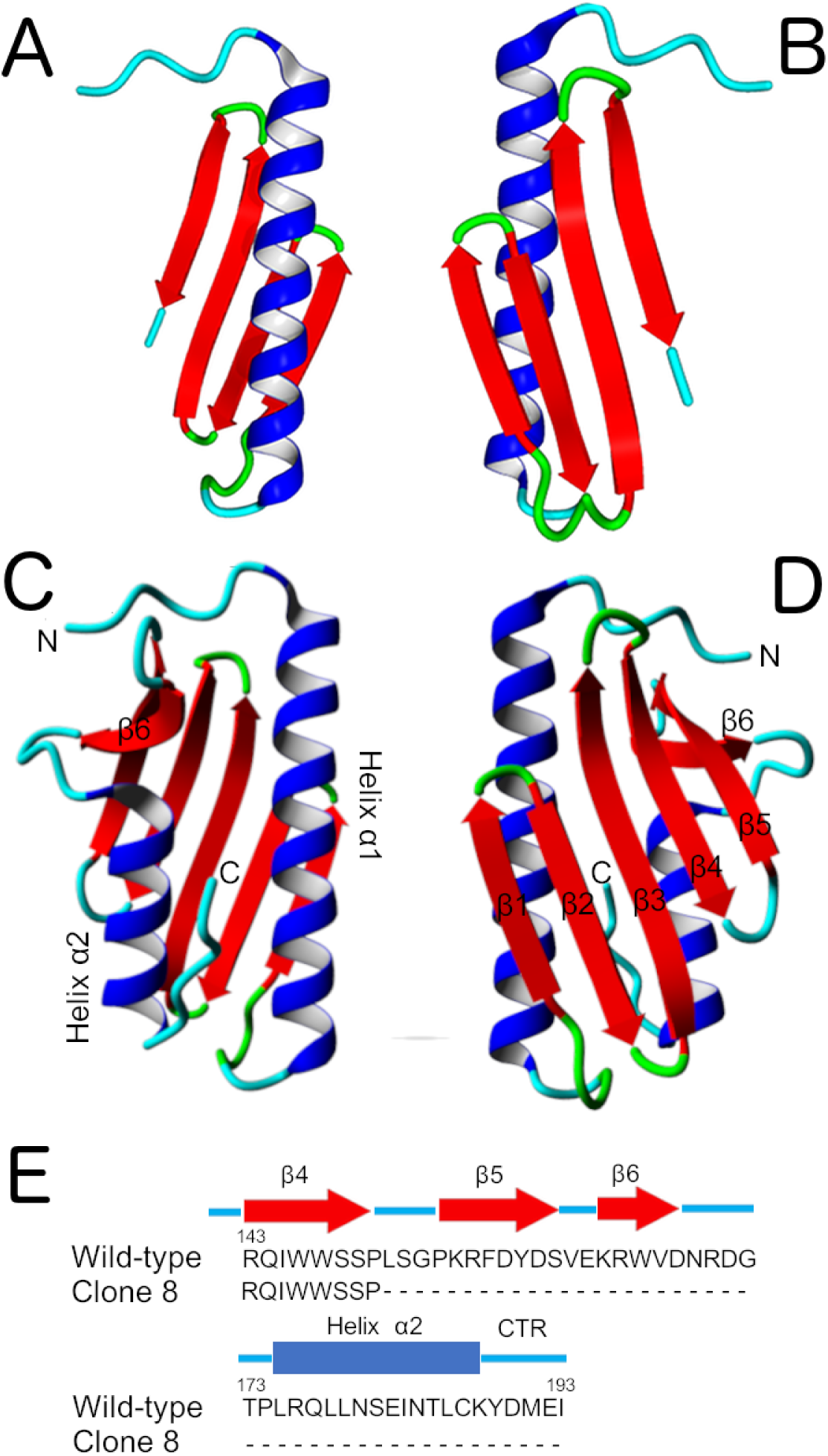
The Alterations of the Frataxin Structure. (A) and (B) Two views of the truncated frataxin DdFXN_81-150 (clone 8). The topology shown is only for visualization purposes. (C) and (D) Two views of the wild-type DdFXN model. DdFXN models were constructed using AlphaFold2 (*52*). AlphaFold formed a disulfide bond between Cys112 and Cys187, whereas other programs like the Swiss-model did not establish the -S-S- bond. (E) Amino acid sequence truncation predicted from the DNA sequences for **clone 8** (**Figure S2** shows the complete sequence alignment).

We studied the molecular motions of this fragment by molecular dynamics simulations. Even though the global conformation of the fragment persisted during 100- 200ns MDs simulations, significantly higher RMSD values and atomic fluctuations were observed, compared to the wild-type (**Figures 4A and B**). Alpha helix 1 presented high distortions, establishing non-native contacts with the beta sheet (**Figures 4E and F**) and higher internal motions compared to the wild-type frataxin, as judged by the fluctuations.

**Figure 4.**
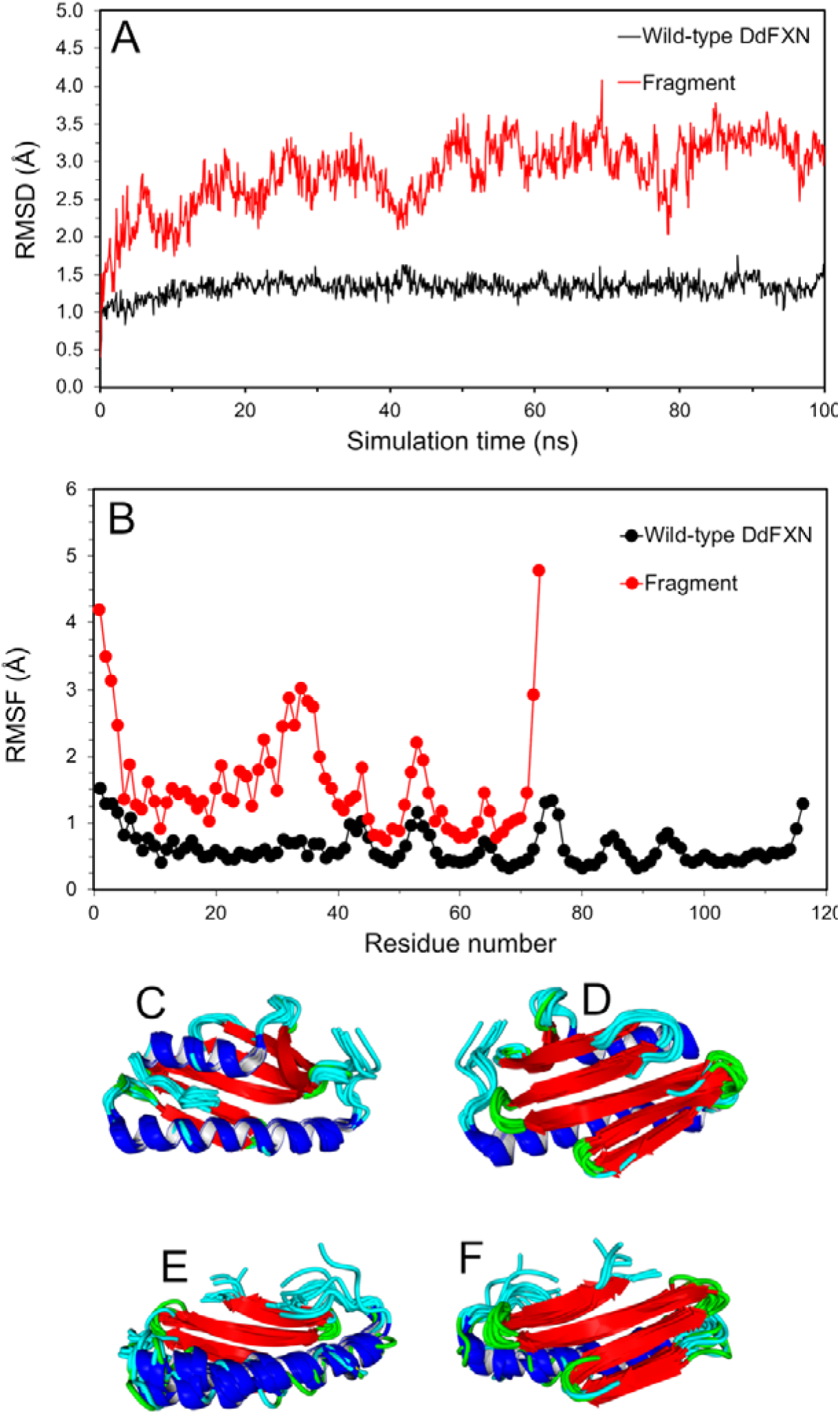
Molecular Dynamics Simulations of the DdFXN Variant. (A) Root-mean-square deviation (RMSD) along the simulations (calculated for the alpha carbon atoms). (B) Root-mean-square fluctuations (RMSF) of the alpha carbon atoms. Snapshots of the wild-type variant (C and D) and the fragment (E and F). Wild-type and clone 8 (fragment) FXN were constructed using AlphaFold2 (*52*). A disulfide bond between Cys112 and Cys187 is formed. For the simulations, three non-native-extra residues in the N-terminal stretch were included: **Met78-Gly79-Ser80-**Pro81-Ile82-Ser83, as for the recombinant proteins produced in our laboratory.

All this suggests that the fragment should be highly dynamic and more likely easily degraded in a cellular environment. However, we cannot rule out that the fragment might acquire a frataxin-like secondary structure and even some kind of packing, although residual activity is highly unlikely.

AlphaFold2 predictions suggested a conformation for a putative dimeric fold of the fragment, in which the predicted accessible apolar surface for each monomer is highly reduced by the interaction, and the remanent Cys residue (Cys112) might establish an intermolecular disulfide bond stabilizing the hypothetical dimeric structure (**Figure S3**). Experiments will be done to evaluate this possibility.

### Edited Clone 8 Presents Undetectable Frataxin Expression Levels

First, we analyzed the expression levels of frataxin in the wild-type AX2 and the edited clone 8 strains. Two different nanobodies that detected the recombinant DdFXN in Western blotting were used to study the frataxin expression in total *D. discoideum* cell lysates (**Figure 5**).

**Figure 5.**
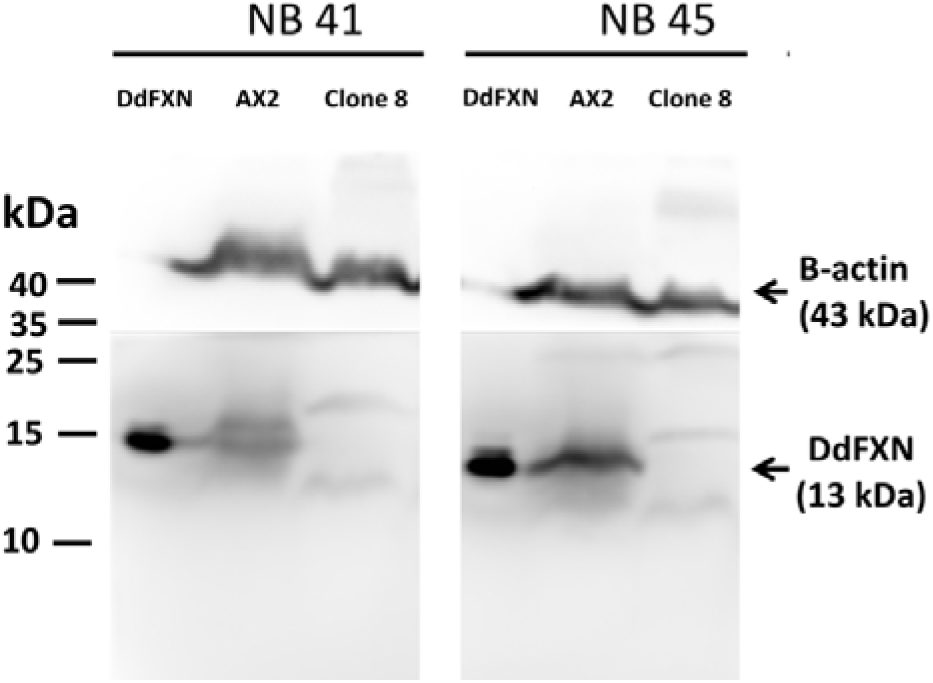
*Analysis of Frataxin Expression by Western Blotting*. Total cell lysates corresponding to AX2 and clone 8 were analyzed by Western blotting using two different nanobodies that carry a 6xHis-tag (NB41 and NB45) raised against DdFXN and a HRP–conjugated secondary antibody antiHis tag (lower panel) and anti ß-actin as loading control (upper panel).

As it can be seen in **Figure 5**, the cell lysate corresponding to the wild-type AX2 undoubtedly showed detectable levels of frataxin. On the other hand, in clone 8 a band corresponding to frataxin or to a frataxin fragment was not detectable. The absence of a detectable signal in clone 8 was not due to the inability of nanobodies to recognize the mutated versions since when the truncated version was heterologous expressed in bacteria, it was readily detected by both nanobodies (**Figure S4**). This result indicates the virtual absence of frataxin in the CRISPR edited cells (**Figure 5**).

### A D. discoideum Clone Lacking Frataxin Has Severe Defective Growth

As an initial global analysis on the effects of altering frataxin functionality in *D. discoideum* biology, we evaluated the growth of the frataxin-deficient clone in the HL5 rich medium and on bacteria. As it is shown in **Figure 6 and Table 3**, clone 8 presents a growth defect in both conditions. The wild-type strain AX2 exhibited a generation time of 9.5 ± 2.2 h, which is in accordance with the literature (*57*). On the other hand, the edited clone 8 carrying the truncated frataxin grew at a lower rate (14.9 ± 2.2 h).

**Figure 6.**
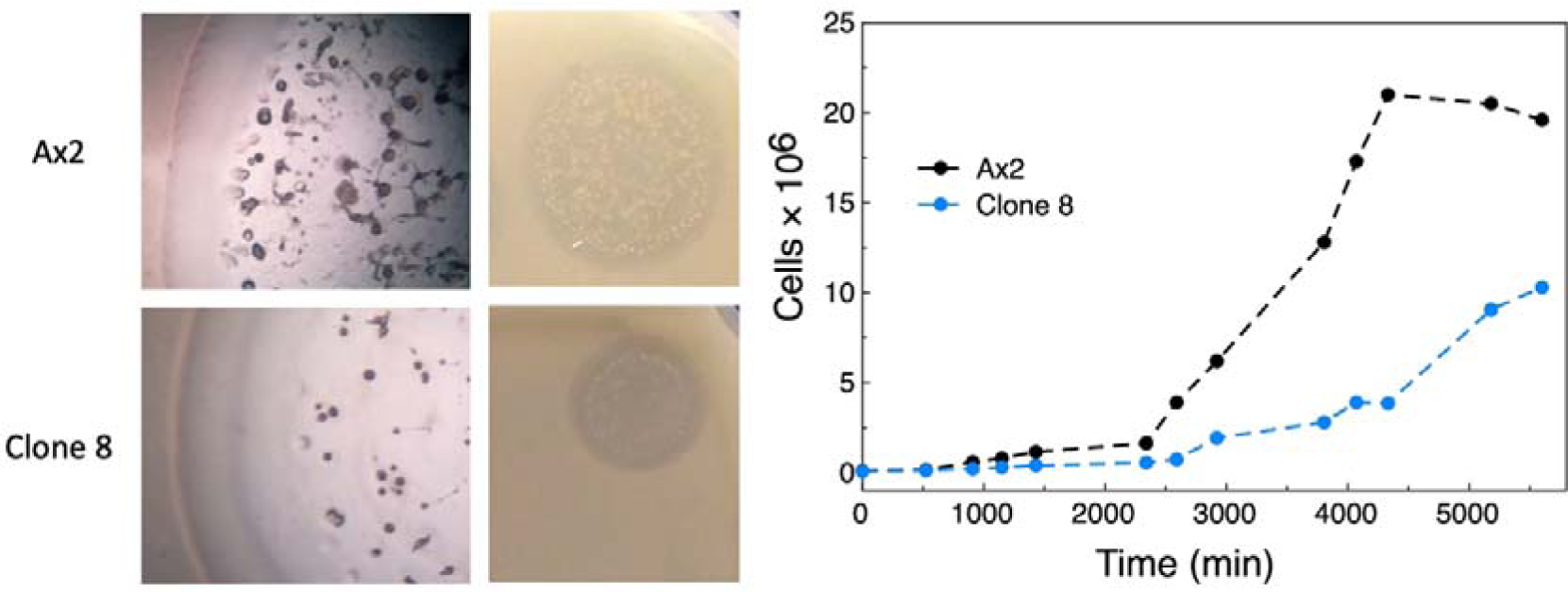
*D. discoideum* Growth. Ax2 and clone 8 were grown on bacterial lawn (left) or in HL5 culture medium (right) starting from a ∼1×10^5^ amoeba cells mL^-1^ inoculum. The cell cultures were grown during a week, and small aliquots were regularly taken for cell counting using a Neubauer chamber. Three independent experiments were carried out; a representative experiment is shown.

**Table 3.** Generation Time Summary.

| Strain | Ax2 | Clone 8 | Ax2 (empty) | Ax2 (G122V) | Ax2 (FXN wt) | Clone 8 (empty) | Clone 8 (G122V) | Clone 8 (FXN wt) |
| --- | --- | --- | --- | --- | --- | --- | --- | --- |
| Generation Time (h) | $9.5 \pm 2.2$ | $14.9 \pm 2.2$ | $12.6 \pm 2.6$ | $13.2 \pm 2.8$ | $12.7 \pm 2.3$ | $19.3 \pm 2.4$ | $12.8 \pm 2.7$ | $13.2 \pm 2.8$ |

### A D. discoideum Clone Lacking Frataxin Presents a Strong Decrease in Fe-S Cluster Dependent Enzymatic Activities

Given that the typical phenotype of a reduction in frataxin functionality is a decrease in Fe-S cluster dependent enzymatic activities, and that this is a feature shared among the FA models, we studied the Krebs cycle enzymes aconitase (ACO) and succinate dehydrogenase (SDH, the Complex II from the respiratory chain). We compared the activity of these mitochondrial enzymes in cellular extracts of strains AX2 and clone 8 (**Figure 7**).

**Figure 7.**
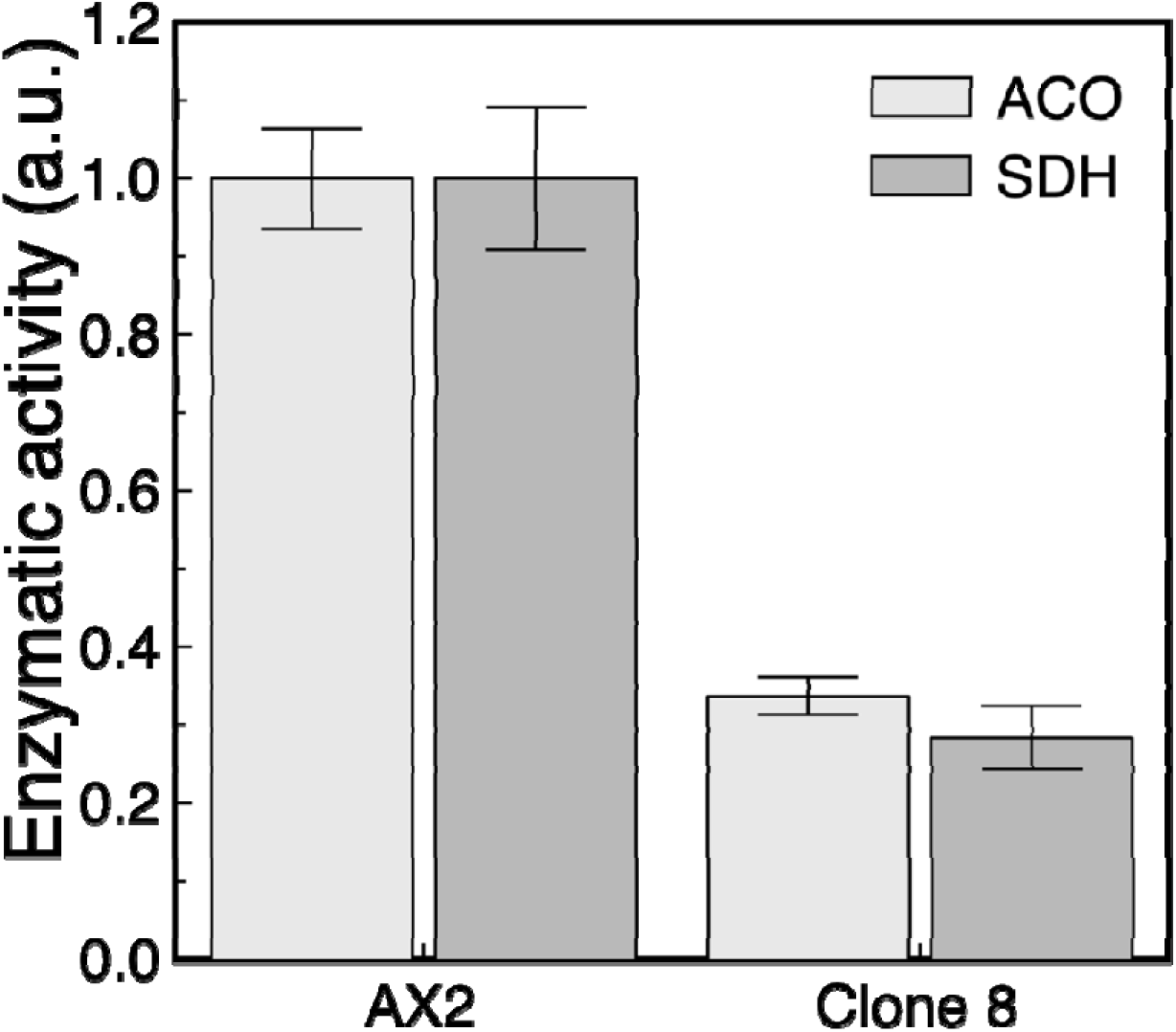
Effect of Frataxin Alteration on Fe-S Cluster Dependent Enzymatic Activities. Aconitase (ACO) and succinate dehydrogenase (SDH) activities were assayed on total lysate extract of *D. discoideum* cells from the AX2 strain and clone 8. Three independent experiments were carried out. The activity measurements are relative to the total protein quantified by triplicate in each extract by the Bradford method.

Clones 8 exhibited a significant reduction of the enzymatic activities by comparison to the wild-type AX2. The activity decreased to 30-40% of the level observed in the wild-type; these values are comparable to those obtained in other cell models deficient in frataxin, where a ∼50% reduction of ACO and SDH activities were detected (*46, 58*).

### A D. discoideum Clone Lacking Frataxin Presents Higher Sensitivity to Oxidative Stress

Another hallmark of FA disease is an increased sensitivity to oxidative stress. The evidence provided by several organism models supports the idea that the deficiency in frataxin function causes a deregulation in antioxidant response (*42, 59–64*). In a similar fashion, one can think that clone 8, which exhibits a significant lower growth rate, may show an inefficient handling of reactive oxygen species (ROS), with an ineffective elimination of ROS by the antioxidant system. To study this issue, we carried out a multi-well plate assay and the effect on cellular viability of a treatment with H_2_O_2_ was determined. We observed a higher impact of hydroperoxide on clone 8 cells compared to the wild-type AX2 cells (**Figure 8**).

**Figure 8.**
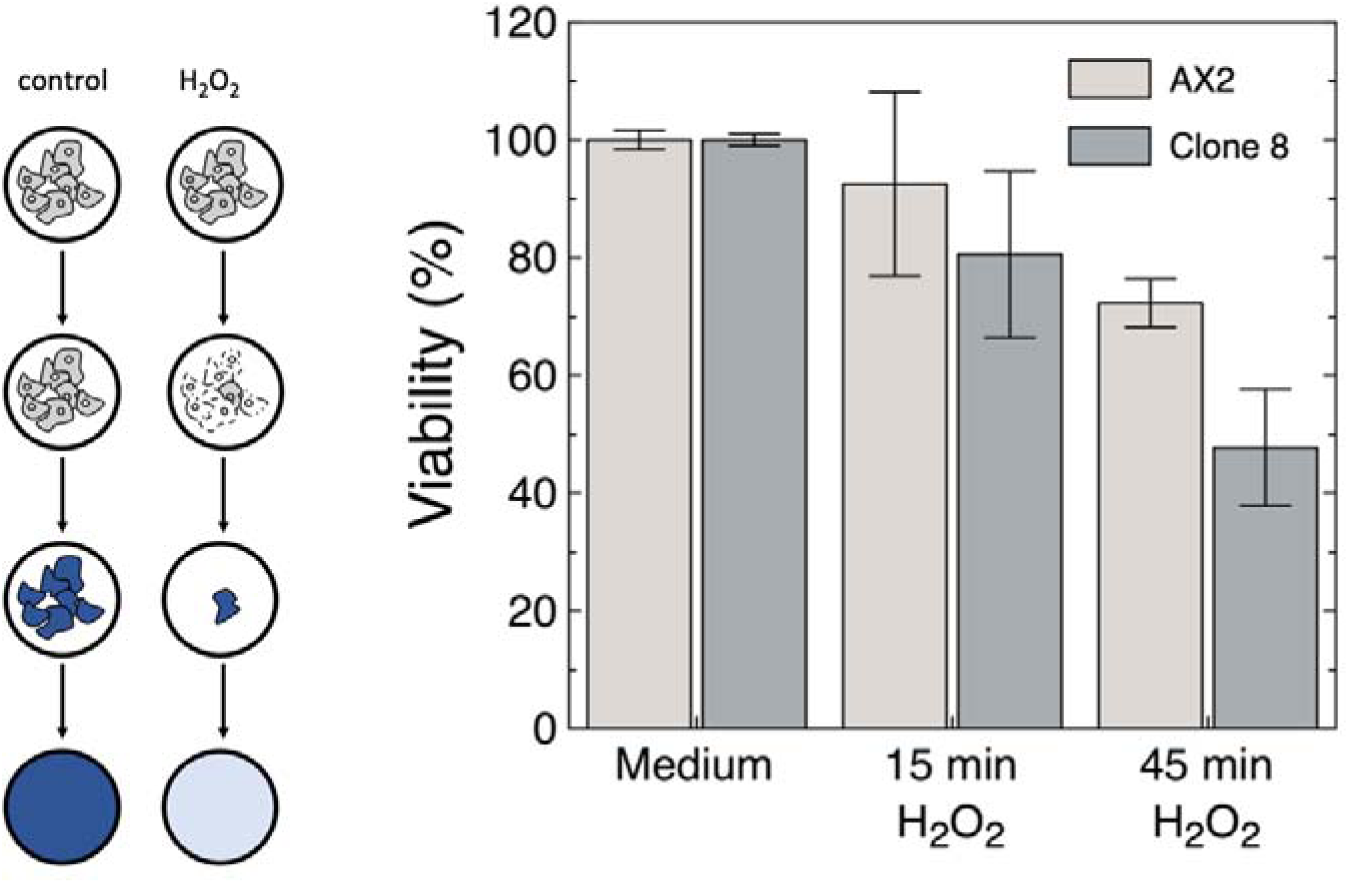
Sensitivity to Oxidative Stress. For determining the viability of cultured cells, a Crystal Violet assay was carried out. The differential effects of H_2_O_2_ on clone 8 cells compared to the wild-type AX2 cells was evaluated. A treatment of frataxin-deficient amoeba with hydroperoxide resulted in a significantly increase of the cell detachment from the multi-well plate surface.

### A D. discoideum Clone Lacking Frataxin Presents Defects in Multicellular Development Progress

To continue with the exploration of the frataxin deficiency effects on *D. discoideum* biology, we monitor the progression of the multicellular development program in the frataxin-deficient clone 8 (**Figure 9**). As mentioned previously, *D. discoideum* grows as isolated ameboid cells that divide mitotically in a nutrient rich culture medium, or in the presence of bacterial lawn. On the other hand, starvation induces a developmental program, the result of which is the formation of a fruiting body bearing spores that disseminate and germinate when nutrients become available. This program involves chemotaxis, cell differentiation, morphogenetic movements, and the integration of internal and external signals. In amoeba with mitochondrial deficiency, as a compensatory mechanism for an energy deficit, there is a chronic increase in AMPK activity (AMP-activated protein kinase) that maintains mitochondrial mass and ATP levels at normal levels; however, it affects other processes that require energy, such as growth, multicellular development, phototaxis and chemotaxis (*65*). Furthermore, previous reports of *D. discoideum* with mitochondrial disease show that mitochondria defective cells present poor growth both in fluid and in bacterial grasses, and also alterations in multicellular development (*66*).

**Figure 9.**
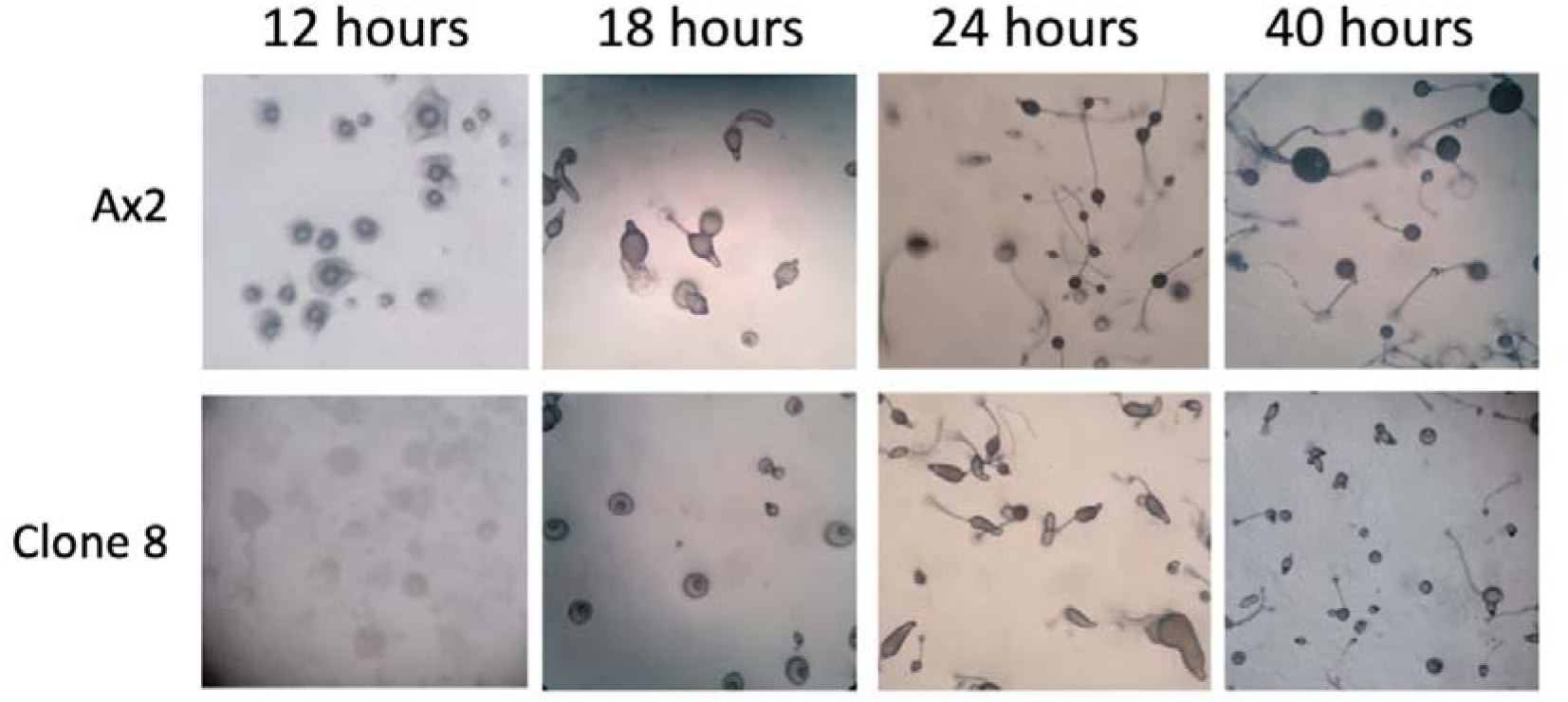
Starvation-Triggered Multicellular Development Progression. Exponentially growing cells of Ax2 (top) and clone 8 (bottom) were seeded in non-nutrient agar plates to induce multicellular development and then incubated at 22 °C. Pictures were taken at different times (12, 18, 24 and 40 hours). A representative picture of each timepoint is shown.

As it can be seen in **Figure 9**, clone 8 presents an altered development; after 18 h of starvation, whereas the wild-type AX2 already exhibited mature fruiting bodies, clone 8 presented immature stages (tipped mound). This significant delay seems to start from the onset of development as it was already evident after 12 h of starvation; at this point, the wild-type strain reached the mound stage while the clone was only at an early loose aggregate stage.

### Constitutive Expression of Frataxin Wild-Type Rescues All Phenotypes in Clone 8

To undoubtedly establish that all defects reported in clone 8 were due to the lack of functional frataxin, we expressed an intron-less, C-terminal flag-tagged wild-type frataxin (DdFXNwt) version from a constitutive promoter.

We then evaluated the capacity of the wild-type DdFXN to rescue the phenotypes reported above. As it can be seen, both growth (**Table 3**) and development (**Figure 10**) of clone 8 expressing DdFXNwt were not significantly different to the wild-type strain AX2 transformed with an empty vector or over-expressing DdFXNwt. On the other hand, clone 8 transformed with an empty vector recapitulates the reported alterations: an increased generation time (**Table 3**), decreased aconitase and succinate dehydrogenase enzymatic activities (**Figure 11B**) and delayed development (**Figure 10**).

**Figure 10.**
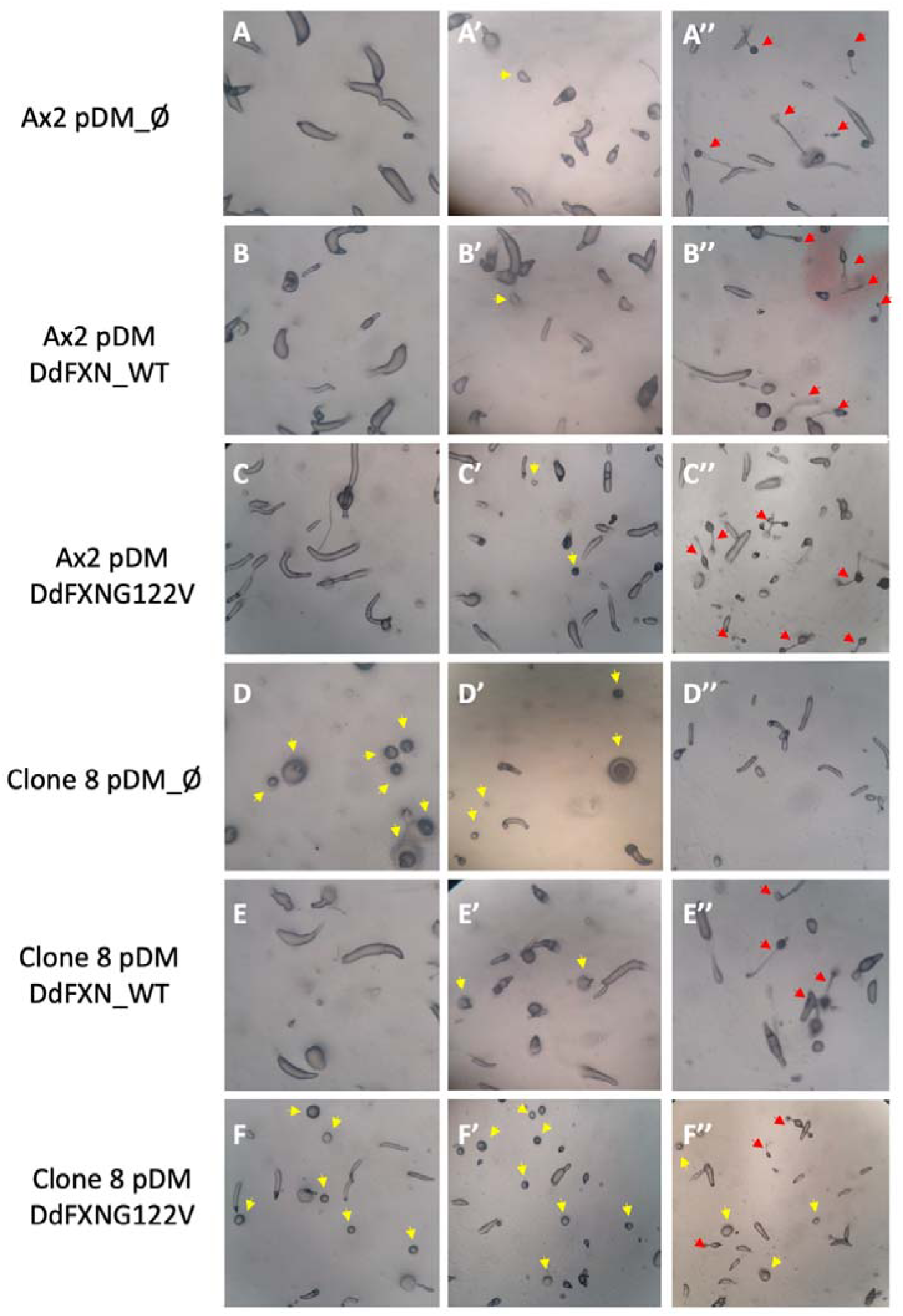
Starvation-Triggered Multicellular Development Progression. Exponentially growing cells of each strain were seeded in non-nutrient agar plates to induce multicellular development and incubated at 22 °C. Representative pictures after 14 hours (left column), 16 hours (middle column) and 20 hours (right column) of development are shown. Yellow arrows and red arrows mark immature mound/tipped mound and early culminants/fruiting bodies.

**Figure 11.**
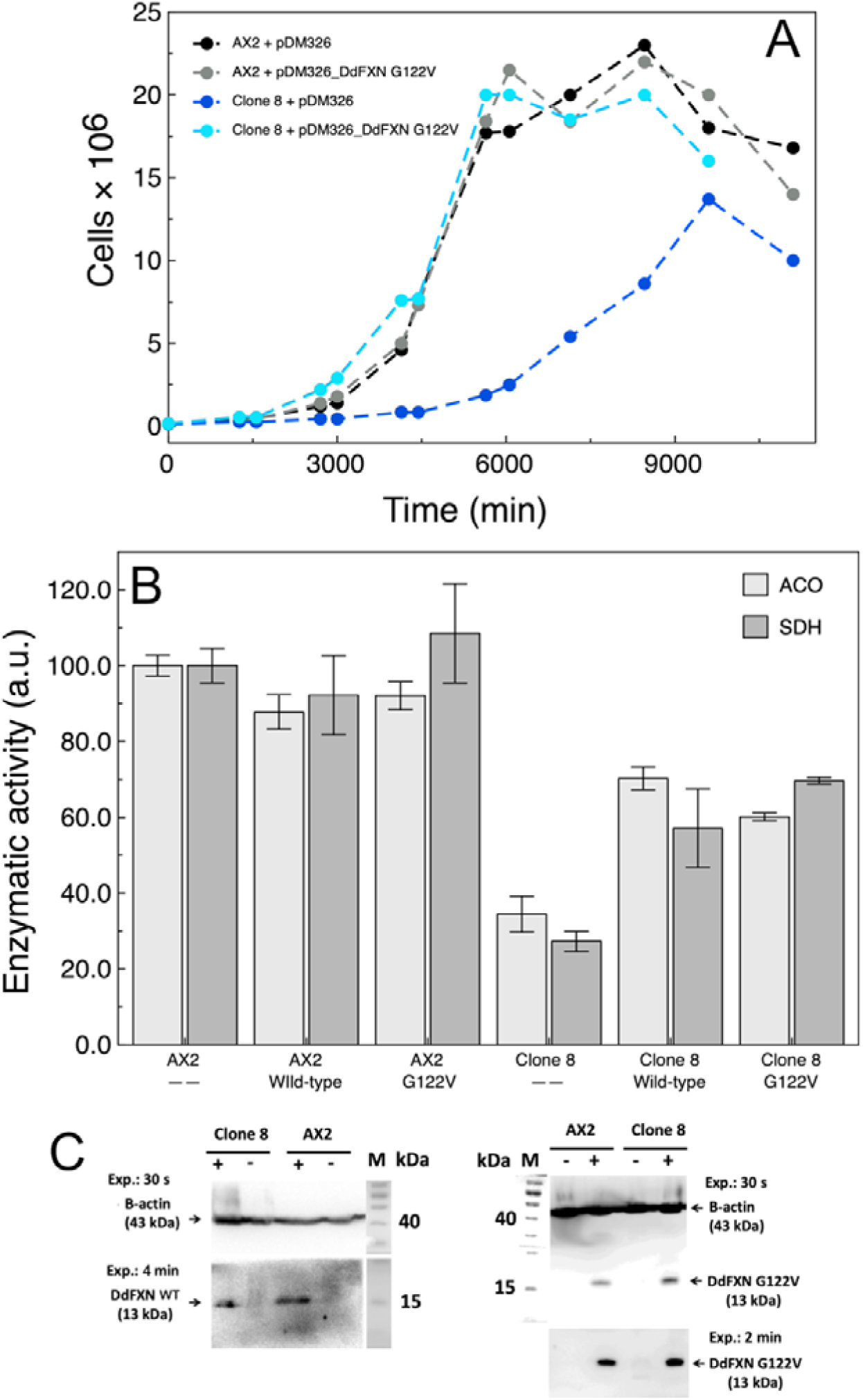
The Expression Frataxin Variant G122V Rescued Clone 8. (A) Growth rate and (B) aconitase and succinate dehydrogenase enzymatic activities of *D. discoideum* AX2 and **clone 8** cells transformed with pDM326, pDM_DdFXNwt or pDM_DdFXN G122V encoding the precursor variant of DdFXN G122V. Growth was carried out at 22 °C and 180 rpm in HL5 culture medium, starting from a 1×10^5^ amoeba cells mL^-1^ inoculum. The cell cultures were grown during a week, and small aliquots were regularly taken for cell counting using a Neubauer chamber. The antibiotic blasticidin was present in culture medium to maintain the pDM326 plasmid in *D. discoideum* cells. For activity assays, the activities were corrected by the total protein mass, measured by Bradford. (C) Western blotting analysis of the expression of DdFXN wild-type (WT) or the DdFXN G122V variant from plasmid pDM326. For G122V, upper and lower panels correspond to 30 s and 2min exposition, respectively. The + and – symbols indicate transformation of *D. discoideum* with the corresponding plasmid without a frataxin gene insert (the wild-type or the DdFXN G122V variant). The detection of the recombinant variant was carried out by using an anti-FLAG monoclonal antibody that recognizes the FLAG sequence located in the C-terminal stretch of the plasmidic frataxin.

Interestingly, the rescue of the decrease in the iron-sulfur cluster dependent enzymatic activities was not complete (**Figure 11B**). The expression of DdFXNwt from the plasmid was corroborated by Western blotting analysis, using an anti-FLAG monoclonal antibody (**Figure 11C**). As expected, no expression was detected when the transformation was carried out with an empty vector.

### Constitutive Expression of Frataxin G122V in Clone 8 Fully Rescues Growth Defect but Produces Only Partial Recovery of Developmental Impairment

Finally, we wondered whether a high expression of a FA frataxin variant can rescue the wild-type features of *D. discoideum* cells. We transformed clone 8 cells with plasmid pDM326-DdFXN-G122V, which encodes the precursor form of G122V frataxin variant. This variant carries a mutation that produces a very specific and less aggressive FA phenotype in humans (G130V). It is worthy of note that clone 8 cells transformed with pDM326-DdFXN-G122V recovered the growth rate of wild-type AX2 (**Table 3**, **Figure 11A**).

We then evaluated whether the G122V frataxin might be able to rescue the defects in multicellular development observed for clone 8. Although a small improvement was detected when these cells were transformed with pDM326-DdFXN- G122V, compared with the same cells transformed with empty pDM326, development was not fully rescued.

At 20 h after starvation, we were able to still see immature structures in clone 8 overexpressing the G122V variant while none were observed when complemented with the wild-type frataxin (**Figure 10**).

We also evaluated the rescue capacity of DdFXN-G122V on the decrease of enzymatic activities (**Figure 11B**); although rescue was not total, the enzymatic levels reached were like the values obtained when DdFXNwt was expressed.

These results suggest that, because of the specific features of this variant, the expression of DdFXN-G122V cannot fully compensate all deficiencies lacking in the wild-type frataxin. This fact also suggests different activities, which are finely tuned by frataxin structure and dynamics, or different controls over the metabolic pathways.

## Discussion and Conclusions

Friedreich’s Ataxia is a rare disease caused by a disfunction of frataxin, a mitochondrial protein, coded in the nucleus. Frataxin is a key component of the mitochondrial supercomplex responsible for Fe-S cluster assembly. Fe-S cluster dependent activities are essential for the homeostasis of the cells. There is a high diversity of cell functions that need the correct insertion of the Fe-S cluster into protein structure. The energetics of the cell, which is commanded by the mitochondrial metabolism, directly depends on Fe-S clusters because clusters are crucial in electron transport reactions. Thus, the alteration of Fe-S cluster biosynthesis affects Complex I, II and III. But Fe-S are also essential for other enzymatic activities like lipoic acid synthesis, necessary for pyruvate dehydrogenase activity or substrate binding in mitochondrial aconitase (Krebs cycle); ultimately, given that pyruvate dehydrogenase complex, Krebs cycle, and the electron transport chain, are all metabolic pathways highly coupled to ATP synthesis, the alteration of the Fe-S cluster assembly results in the impairment of numerous ATP-dependent cellular processes.

In previous research we showed that *D. discoideum* has a complete pathway involved in Fe-S cluster assembly and transferring, as found in mammalian cells. Our inferences from protein sequence conservation and structure models suggested that some proteins (human and *D. discoideum*) may be exchangeable. In fact, the FXN residues that when mutated result in FA are practically fully conserved between human and *D. discoideum*. In this study, we observed that DdFXN can bind to the human supercomplex NIAU with a high affinity, similarly to the human FXN variant. Additionally, it was able to activate L-Cys desulfurase function. This is indicative of the role of protein-protein interaction surfaces and the high conservation score for the residues involved in these interactions. This highlights the potential of *D. discoideum* as a model system for FA.

We then showed that frataxin locus can be easily edited in *D. discoideum*. Taking advantage of the haploid context of this amoeba, we constructed a completely frataxin-deficient strain. In the edited *D. discoideum* clone 8, studied in this paper, a premature STOP codon occurred. The frataxin fragment encoded in the genomic DNA seemed to be extremely unstable as judged by its absence in clone 8 amoeba cellular lysates, a result that matches with molecular dynamics simulations, which suggests a highly mobile protein backbone compared to the wild-type DdFXN. This strain presents a significant decrease of the Fe-S cluster dependent enzymatic functions and an altered phenotype. Reduced aconitase and succinate dehydrogenase activities, a decrease in growth rate, and higher sensitivity to oxidative stress were consistently observed, being all these landmarks of frataxin deficiency in other Friedreich’s Ataxia cellular and organism models. Furthermore, we explored other processes that have been linked to mitochondrial disfunction in *D. discoideum* and found defects during multicellular development. Remarkably, we were able to rescue phenotype alterations with the constitutive expression of wild-type DdFXN. The expression of the wild-type protein was able to fully rescue growth and multicellular behavior alterations while the decreased enzymatic activities were only partially rescued. There are a few conceivable, not mutually exclusive, explanations for this result. The frataxin expression levels obtained when expressed from plasmids are different than endogenous. Furthermore, given that frataxin is imported into the mitochondrial matrix, the higher expression levels of the protein might also affect the importing and processing machinery, although no effect was observed when a constitutive expression was carried out in the wild-type AX2 strain. On the other hand, it is also worth noting that the DdFXN sequence used for protein expression from plasmids was engineered carrying a C-terminal FLAG tag for protein identity assignment, which could affect some roles of frataxin but not others. The fact that the FLAG sequence includes a highly negatively charged stretch of residues, makes feasible that it alters in some degree frataxin interaction with the *D. discoideum* NIAU supercomplex. Experiments with untagged versions would help to test this hypothesis.

We then used this set-up to assess the rescuing capacity of DdFXN G122V, a frataxin version that mimics a pathogenic variant found in Friedreich’s Ataxia patients (i.e., G122V in *D. discoideum* corresponds to G130V in humans). This variant, when it is constitutively expressed, fully rescues the defects on growth but only marginally rescues the defects on multicellular development. This underlies the fact that DdFXN G122V is not merely a less stable frataxin variant, because increased expression levels should bypass the defect. So it may be that other features of the protein affected in G122V are important for specific roles played by frataxin. Hence, clone 8 can help to dissect the different functions of FXN within the cell. On the other hand, the defects of enzymatic activities were rescued to the same levels as in the case of the expression of the wild-type C-terminal tagged DdFXN.

In this context, this new biological model offers a wide range of options to easily explore diverse phenotypes occurring in FA; this makes *D. discoideum* a very attractive approach for studying, in a straightforward manner, the effect that diversity of FA variants has on the cellular metabolism. Moreover, this model may help to understand whether frataxin works as a bottleneck for some processes, whereas in other processes it has lower flux control, at least under the experimental conditions assayed.

In summary, we have generated a system where we can easily monitor the effects of a lack of frataxin activity; this opens the door to developing drug or treatment screenings that would help to design and/or evaluate therapeutical strategies. Moreover, this biological model offers a wide range of possibilities to easily explore diverse phenotypes in FA. To complete the list of desirable attributes for a disease model organism, it has been proved that frataxin locus can be easily edited, which enables the generation of specific strains that express the variant of interest from an endogenous locus.

## Acknowledgments

We are grateful to CONICET and Universidad de Buenos Aires. We especially thank Fundación Ciencias Exactas y Naturales (FUNDACEN) and ALAPA. We are also greateful to Dr. Guillermo D. Alonso for its help with DdFXN cloning and general support of the project.

## Abbreviations

ACP: acyl carrier protein
Cas9: CRISPR associated protein 9
CD: circular dichroism
CTR: C-terminal region
CRISPR: clustered regularly interspaced short palindromic repeats
DLS: dynamic light scattering
Fe-S: iron-sulfur
FA: Friedreich’s Ataxia
FXN: frataxin
DdFXN: *D. discoideum* frataxin
DTT: dithiothreitol
HPLC: high-performance liquid chromatography
ISCU: iron-sulfur cluster assembly enzyme
ISD11: NFS1 interacting protein
NFS1: mitochondrial L- cysteine desulfurase enzyme
NMR: nuclear magnetic resonance
PAGE: polyacrylamide gel electrophoresis
PDB: Protein Data Bank
SDS: sodium dodecyl sulfate
SEC: size exclusion chromatography

## References

1. L. Fets, R. Kay, F. Velazquez, Dictyostelium. Current Biology. 20, R1008–R1010 (2010).

2. J. M. Goldberg, G. Manning, A. Liu, P. Fey, K. E. Pilcher, Y. Xu, J. L. Smith, The Dictyostelium Kinome—Analysis of the Protein Kinases from a Simple Model Organism. PLoS Genet. 2, e38 (2006).

3. J. D. Dunn, C. Bosmani, C. Barisch, L. Raykov, L. H. Lefrançois, E. Cardenal- Muñoz, A. T. López-Jiménez, T. Soldati, Eat Prey, Live: Dictyostelium discoideum As a Model for Cell-Autonomous Defenses. Front Immunol. 8, 1906 (2018).

4. J. Martín-González, J.-F. Montero-Bullón, J. Lacal, Dictyostelium discoideum as a non-mammalian biomedical model. Microb Biotechnol. 14, 111–125 (2021).

5. H. N. Haver, K. M. Scaglione, Dictyostelium discoideum as a Model for Investigating Neurodegenerative Diseases. Front Cell Neurosci. 15 (2021), doi:10.3389/FNCEL.2021.759532.

6. J. Schaf, J. Damstra-Oddy, R. S. B. Williams, Dictyostelium discoideum as a pharmacological model system to study the mechanisms of medicinal drugs and natural products. International Journal of Developmental Biology. 63, 541–550 (2019).

7. K. Augustin, A. Khabbush, S. Williams, S. Eaton, M. Orford, J. H. Cross, S. J. R. Heales, M. C. Walker, R. S. B. Williams, Mechanisms of action for the medium-chain triglyceride ketogenic diet in neurological and metabolic disorders. Lancet Neurol. 17, 84–93 (2018).

8. R. H. Kessin, Dictyostelium (Cambridge University Press, Cambridge, 2001; http://ebooks.cambridge.org/ref/id/CBO9780511525315).

9. G. Bloomfield, Sex and macrocyst formation in Dictyostelium. International Journal of Developmental Biology. 63, 439–446 (2019).

10. K. Yamashita, H. Iriki, Y. Kamimura, T. Muramoto, CRISPR Toolbox for Genome Editing in Dictyostelium. Front Cell Dev Biol. 9 (2021), doi:10.3389/FCELL.2021.721630.

11. T. Ogasawara, J. Watanabe, R. Adachi, Y. Ono, Y. Kamimura, T. Muramoto, CRISPR/Cas9-based genome-wide screening of Dictyostelium. Scientific Reports 2022 12:1. 12, 1–13 (2022).

12. T. Muramoto, H. Iriki, J. Watanabe, T. Kawata, Recent Advances in CRISPR/Cas9-Mediated Genome Editing in Dictyostelium. Cells. 8, 46 (2019).

13. G. P. Otto, M. Cocorocchio, L. Munoz, R. A. Tyson, T. Bretschneider, R. S. B. Williams, “Employing Dictyostelium as an advantageous 3Rs model for pharmacogenetic research” in Methods in Molecular Biology (Humana Press Inc., 2016; https://pubmed.ncbi.nlm.nih.gov/27271898/), vol. 1407, pp. 123–130.

14. A. Tornero-Écija, L. C. Tábara, M. Bueno-Arribas, L. Antón-Esteban, C. Navarro-Gómez, I. Sánchez, O. Vincent, R. Escalante, A Dictyostelium model for BPAN disease reveals a functional relationship between the WDR45/WIPI4 homolog Wdr45l and Vmp1 in the regulation of autophagy-associated PtdIns3P and ER stress. Autophagy. 18, 661–677 (2022).

15. J. Rodriguez-Centeno, R. Perona, L. Sastre, Dyskerin Mutations Present in Dyskeratosis Congenita Patients Increase Oxidative Stress and DNA Damage Signalling in Dictyostelium Discoideum. Cells 2019, Vol. 8, Page 1406. 8, 1406 (2019).

16. A. C. Adam, C. Bornhövd, H. Prokisch, W. Neupert, K. Hell, The Nfs1 interacting protein Isd11 has an essential role in Fe/S cluster biogenesis in mitochondria. EMBO Journal. 25, 174–183 (2006).

17. S. C. Lim, M. Friemel, J. E. Marum, E. J. Tucker, D. L. Bruno, L. G. Riley, J. Christodoulou, E. P. Kirk, A. Boneh, C. M. DeGennaro, M. Springer, V. K. Mootha, T. A. Rouault, S. Leimkühler, D. R. Thorburn, A. G. Compton, Mutations in LYRM4, encoding iron-sulfur cluster biogenesis factor ISD11, cause deficiency of multiple respiratory chain complexes. Hum Mol Genet. 22, 4460–4473 (2013).

18. L. Böttinger, C. U. Martensson, J. Song, N. Zufall, N. Wiedemann, T. Becker, Respiratory chain supercomplexes associate with the cysteine desulfurase complex of the iron-sulfur cluster assembly machinery. Mol Biol Cell. 29, 776– 785 (2018).

19. J. K. Barton, R. M. B. Silva, E. O’Brien, Redox chemistry in the genome: Emergence of the [4fe4s] cofactor in repair and replication. Annu Rev Biochem. 88, 163–190 (2019).

20. A. K. Pandey, J. Pain, A. Dancis, D. Pain, Mitochondria export iron-sulfur and sulfur intermediates to the cytoplasm for iron-sulfur cluster assembly and tRNA thiolation in yeast. Journal of Biological Chemistry. 294, 9489–9502 (2019).

21. J. A. Mayr, R. G. Feichtinger, F. Tort, A. Ribes, W. Sperl, Lipoic acid biosynthesis defects. J Inherit Metab Dis. 37, 553–563 (2014).

22. T. A. Richards, M. van der Giezen, Evolution of the Isd11-IscS complex reveals a single alpha-proteobacterial endosymbiosis for all eukaryotes. Mol Biol Evol. 23, 1341–1344 (2006).

23. M. Georgina Herrera, M. Florencia Pignataro, M. Ezequiel Noguera, K. Magalí Cruz, J. Santos, M. G. Herrera, M. F. Pignataro, M. E. Noguera, K. M. Cruz, J. Santos, Rescuing the Rescuer: On the Protein Complex between the Human Mitochondrial Acyl Carrier Protein and ISD11. ACS Chem Biol. 13, 1455–1462 (2018).

24. K. Cai, R. O. Frederick, M. Tonelli, J. L. Markley, Mitochondrial Cysteine Desulfurase and ISD11 Coexpressed in Escherichia coli Yield Complex Containing Acyl Carrier Protein. ACS Chem Biol. 12, 918–921 (2017).

25. M. G. Herrera, M. E. Noguera, K. E. Sewell, W. A. Agudelo Suárez, L. Capece, S. Klinke, J. Santos, Structure of the Human ACP-ISD11 Heterodimer. Biochemistry. 58 (2019), doi:10.1021/acs.biochem.9b00539.

26. N. G. Fox, X. Yu, X. Feng, H. J. Bailey, A. Martelli, J. F. Nabhan, C. Strain-Damerell, C. Bulawa, W. W. Yue, S. Han, Structure of the human frataxin-bound iron-sulfur cluster assembly complex provides insight into its activation mechanism. 10, 2210 (2019).

27. S. Gervason, D. Larkem, A. ben Mansour, T. Botzanowski, C. S. Müller, L. Pecqueur, G. le Pavec, A. Delaunay-Moisan, O. Brun, J. Agramunt, A. Grandas, M. Fontecave, V. Schünemann, S. Cianférani, C. Sizun, M. B. Tolédano, B. D’Autréaux, Physiologically relevant reconstitution of iron-sulfur cluster biosynthesis uncovers persulfide-processing functions of ferredoxin-2 and frataxin. Nat Commun. 10 (2019), doi:10.1038/S41467-019-11470-9.

28. S. M. K. Farhan, J. Wang, J. F. Robinson, P. Lahiry, V. M. Siu, C. Prasad, J. B. Kronick, D. A. Ramsay, C. Anthony Rupar, R. A. Hegele, Exome sequencing identifies nfs1 deficiency in a novel fe-s cluster disease, infantile mitochondrial complex ii/iii deficiency. Mol Genet Genomic Med. 2, 73–80 (2014).

29. F. Mochel, M. A. Knight, W. H. Tong, D. Hernandez, K. Ayyad, T. Taivassalo, P. M. Andersen, A. Singleton, T. A. Rouault, K. H. Fischbeck, R. G. Haller, Splice Mutation in the Iron-Sulfur Cluster Scaffold Protein ISCU Causes Myopathy with Exercise Intolerance. Am J Hum Genet. 82, 652–660 (2008).

30. S. E. Faraj, E. A. Roman, M. Aran, M. Gallo, J. Santos, The alteration of the C- terminal region of human frataxin distorts its structural dynamics and function. FEBS Journal. 281, 3397–3419 (2014).

31. N. Faggianelli, R. Puglisi, L. Veneziano, S. Romano, M. Frontali, T. Vannocci, S. Fortuni, R. Testi, A. Pastore, Analyzing the effects of a G137V mutation in the FXN gene. Front Mol Neurosci. 8, 1–8 (2015).

32. A. R. Correia, C. Pastore, S. Adinolfi, A. Pastore, C. M. Gomes, Dynamics, stability and iron-binding activity of frataxin clinical mutants. FEBS J. 275, 3680–3690 (2008).

33. P. Cavadini, C. Gellera, P. I. Patel, G. Isaya, Human frataxin maintains mitochondrial iron homeostasis in Saccharomyces cerevisiae. Hum Mol Genet. 9, 2523–2530 (2000).

34. D. Doni, L. Passerini, G. Audran, S. R. A. Marque, M. Schulz, J. Santos, P. Costantini, M. Bortolus, D. Carbonera, Effects of Fe2+/Fe3+ Binding to Human Frataxin and Its D122Y Variant, as Revealed by Site-Directed Spin Labeling (SDSL) EPR Complemented by Fluorescence and Circular Dichroism Spectroscopies. International Journal of Molecular Sciences 2020, Vol. 21, Page 9619. 21, 9619 (2020).

35. C. L. Tsai, J. Bridwell-Rabb, D. P. Barondeau, Friedreich’s ataxia variants I154F and W155R diminish frataxin-based activation of the iron-sulfur cluster assembly complex. Biochemistry. 50, 6478 (2011).

36. J. Bridwell-Rabb, A. M. Winn, D. P. Barondeau, Structure - Function analysis of friedreich’s ataxia mutants reveals determinants of frataxin binding and activation of the fe - S assembly complex. Biochemistry. 50, 7265–7274 (2011).

37. F. Saccà, A. Marsili, G. Puorro, A. Antenora, C. Pane, A. Tessa, P. Scoppettuolo, C. Nesti, V. Brescia Morra, G. de Michele, F. M. Santorelli, A. Filla, Clinical use of frataxin measurement in a patient with a novel deletion in the FXN gene. J Neurol. 260, 1116–1121 (2013).

38. D. Poburski, J. B. Boerner, M. Koenig, M. Ristow, R. Thierbach, Time-resolved functional analysis of acute impairment of frataxin expression in an inducible cell model of Friedreich ataxia. Biol Open. 5, 654 (2016).

39. M. Cossée, H. Puccio, A. Gansmuller, H. Koutnikova, A. Dierich, M. LeMeur, K. Fischbeck, P. Dollé, M. Kœnig, Inactivation of the Friedreich ataxia mouse gene leads to early embryonic lethality without iron accumulation. Hum Mol Genet. 9, 1219–1226 (2000).

40. M. v. Busi, M. v. Maliandi, H. Valdez, M. Clemente, E. J. Zabaleta, A. Araya, D. F. Gomez-Casati, Deficiency of Arabidopsis thaliana frataxin alters activity of mitochondrial Fe–S proteins and induces oxidative stress. The Plant Journal. 48, 873–882 (2006).

41. N. Ventura, S. L. Rea, R. Testi, Long-lived C. elegans Mitochondrial mutants as a model for human mitochondrial-associated diseases. Exp Gerontol. 41, 974– 991 (2006).

42. J. v. Llorens, J. A. Navarro, M. J. Martínez-Sebastián, M. K. Baylies, S. Schneuwly, J. A. Botella, M. D. Moltó, Causative role of oxidative stress in a Drosophila model of Friedreich ataxia. The FASEB Journal. 21, 333–344 (2007).

43. P. R. Anderson, K. Kirby, A. J. Hilliker, J. P. Phillips, RNAi-mediated suppression of the mitochondrial iron chaperone, frataxin, in Drosophila. Hum Mol Genet. 14, 3397–3405 (2005).

44. I. Zanella, M. Derosas, M. Corrado, E. Cocco, P. Cavadini, G. Biasiotto, M. Poli, R. Verardi, P. Arosio, The effects of frataxin silencing in HeLa cells are rescued by the expression of human mitochondrial ferritin. Biochimica et Biophysica Acta (BBA) - Molecular Basis of Disease. 1782, 90–98 (2008).

45. C. Lu, G. Cortopassi, Frataxin knockdown causes loss of cytoplasmic iron–sulfur cluster functions, redox alterations and induction of heme transcripts. Arch Biochem Biophys. 457, 111 (2007).

46. N. Calmels, S. Schmucker, M. Wattenhofer-Donzé, A. Martelli, N. Vaucamps, L. Reutenauer, N. Messaddeq, C. Bouton, M. Koenig, H. Puccio, The first cellular models based on frataxin missense mutations that reproduce spontaneously the defects associated with Friedreich ataxia. PLoS One. 4 (2009), doi:10.1371/journal.pone.0006379.

47. F. Lupoli, T. Vannocci, G. Longo, N. Niccolai, A. Pastore, The role of oxidative stress in Friedreich’s ataxia. FEBS Lett. 592, 718–727 (2018).

48. F. Codazzi, A. Hu, M. Rai, S. Donatello, F. S. Scarzella, E. Mangiameli, I. Pelizzoni, F. Grohovaz, M. Pandolfo, Friedreich ataxia-induced pluripotent stem cell-derived neurons show a cellular phenotype that is corrected by a benzamide HDAC inhibitor. Hum Mol Genet. 25 (2016), doi:10.1093/hmg/ddw308.

49. J. Olmos, M. F. Pignataro, A. B. B. dos Santos, M. Bringas, S. Klinke, L. Kamenetzky, F. Velazquez, J. Santos, A highly conserved iron-sulfur cluster assembly machinery between humans and amoeba dictyostelium discoideum: The characterization of frataxin. Int J Mol Sci. 21, 1–25 (2020).

50. R. Sekine, T. Kawata, T. Muramoto, CRISPR/Cas9 mediated targeting of multiple genes in Dictyostelium OPEN. Scientific RepoRts |. 8, 8471 (2018).

51. H. Land, M. S. Humble, YASARA: A Tool to Obtain Structural Guidance in Biocatalytic Investigations. Methods Mol Biol. 1685, 43–67 (2018).

52. R. Evans, M. O’Neill, A. Pritzel, N. Antropova, A. Senior, T. Green, A. Žídek, R. Bates, S. Blackwell, J. Yim, O. Ronneberger, S. Bodenstein, M. Zielinski, A. Bridgland, A. Potapenko, A. Cowie, K. Tunyasuvunakool, R. Jain, E. Clancy, P. Kohli, J. Jumper, D. Hassabis, bioRxiv, in press, doi:10.1101/2021.10.04.463034.

53. J. A. Maier, C. Martinez, K. Kasavajhala, L. Wickstrom, K. E. Hauser, C. Simmerling, ff14SB: Improving the Accuracy of Protein Side Chain and Backbone Parameters from ff99SB. J Chem Theory Comput. 11, 3696–3713 (2015).

54. E. Krieger, G. Vriend, New ways to boost molecular dynamics simulations. J Comput Chem. 36, 996 (2015).

55. M. Feoktistova, P. Geserick, M. Leverkus, Cold Spring Harb Protoc, in press, doi:10.1101/PDB.PROT087379.

56. T. J. Gibson, E. v Koonin, G. Musco, A. Pastore, P. Bork, Friedreich’s ataxia protein: Phylogenetic evidence for mitochondrial dysfunction. Trends Neurosci. 19, 465–468 (1996).

57. J. M. Ashworth, D. J. Watts, Metabolism of the cellular slime mould Dictyostelium discoideum grown in axenic culture. Biochemical Journal. 119, 175 (1970).

58. T. Ast, J. D. Meisel, S. Patra, H. Wang, R. M. H. Grange, S. H. Kim, S. E. Calvo, L. L. Orefice, F. Nagashima, F. Ichinose, W. M. Zapol, G. Ruvkun, D. P. Barondeau, V. K. Mootha, Hypoxia Rescues Frataxin Loss by Restoring Iron Sulfur Cluster Biogenesis. Cell. 177, 1507–1521.e16 (2019).

59. S. Al-Mahdawi, R. M. Pinto, D. Varshney, L. Lawrence, M. B. Lowrie, S. Hughes, Z. Webster, J. Blake, J. M. Cooper, R. King, M. A. Pook, GAA repeat expansion mutation mouse models of Friedreich ataxia exhibit oxidative stress leading to progressive neuronal and cardiac pathology. Genomics. 88, 580–590 (2006).

60. K. Chantrel-Groussard, V. Geromel, H. Puccio, M. Koenig, A. Munnich, A. Rötig, P. Rustin, Disabled early recruitment of antioxidant defenses in Friedreich’s ataxia. Hum Mol Genet. 10, 2061–2067 (2001).

61. C. M. Gomes, R. Santos, Neurodegeneration in Friedreich’s Ataxia: From Defective Frataxin to Oxidative Stress. Oxid Med Cell Longev. 2013 (2013), doi:10.1155/2013/487534.

62. R. Santos, N. Buisson, S. A. B. Knight, A. Dancis, J. M. Camadro, E. Lesuisse, Candida albicans lacking the frataxin homologue: a relevant yeast model for studying the role of frataxin. Mol Microbiol. 54, 507–519 (2004).

63. R. P. Vázquez-Manrique, P. González-Cabo, S. Ros, H. Aziz, H. A. Baylis, F. Palau, Reduction of Caenorhabditis elegans frataxin increases sensitivity to oxidative stress, reduces lifespan, and causes lethality in a mitochondrial complex II mutant. The FASEB Journal. 20, 172–174 (2006).

64. A. Wong, J. Yang, P. Cavadini, C. Gellera, B. Lonnerdal, F. Taroni, G. Cortopassi, The Friedreich’s Ataxia Mutation Confers Cellular Sensitivity to Oxidant Stress Which Is Rescued by Chelators of Iron and Calcium and Inhibitors of Apoptosis. Hum Mol Genet. 8, 425–430 (1999).

65. L. M. Francione, S. J. Annesley, S. Carilla-Latorre, R. Escalante, P. R. Fisher, The Dictyostelium model for mitochondrial disease. Semin Cell Dev Biol. 22, 120–130 (2011).

66. M. Kotsifas, C. Barth, A. de Lozanne, S. T. Lay, P. R. Fisher, Chaperonin 60 and mitochondrial disease in Dictyostelium.

