## Supplemental Figures for "CRISPR/Cas9-Based Edition of Frataxin Gene in *Dictyostelium discoideum* for Friedreich’s Ataxia Disease Modeling"

**Figure S1. Summary of frataxin locus CRISPR/Cas edition in clones - preliminary characterization**

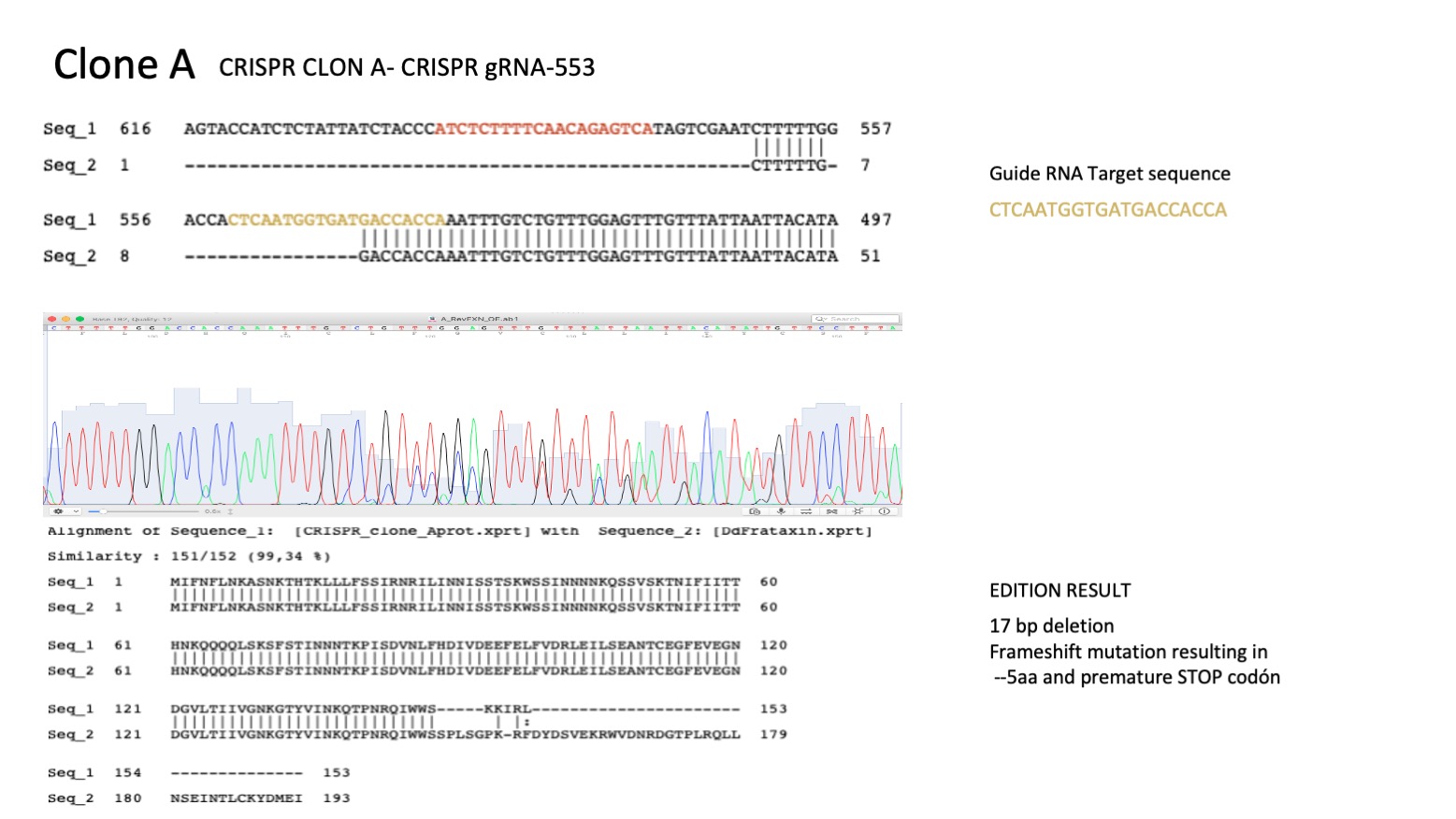

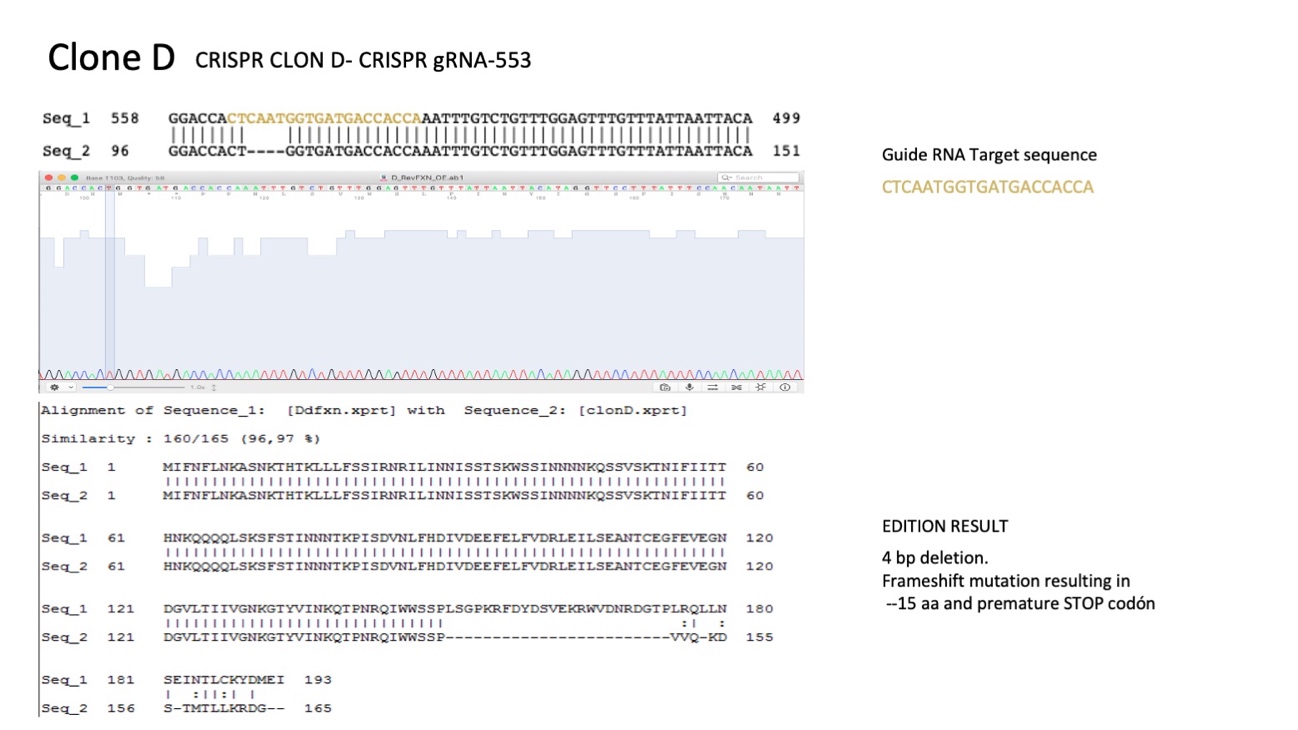

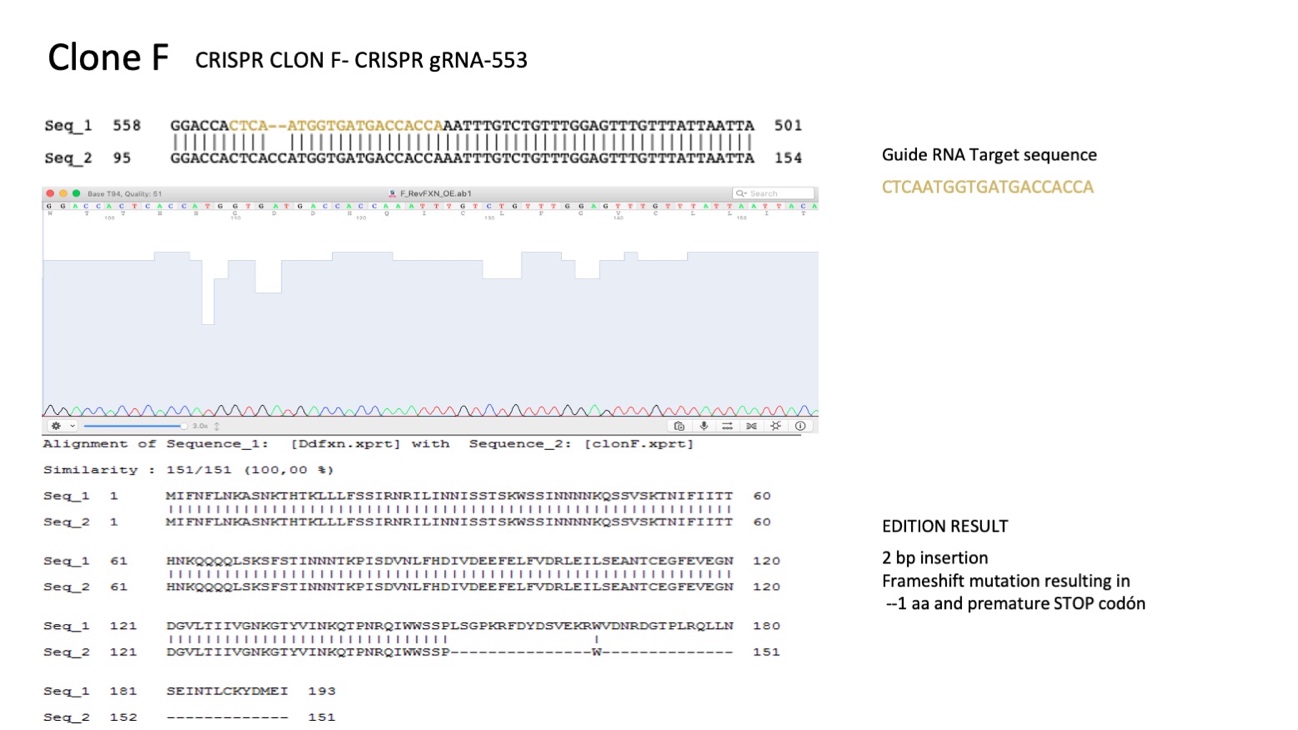

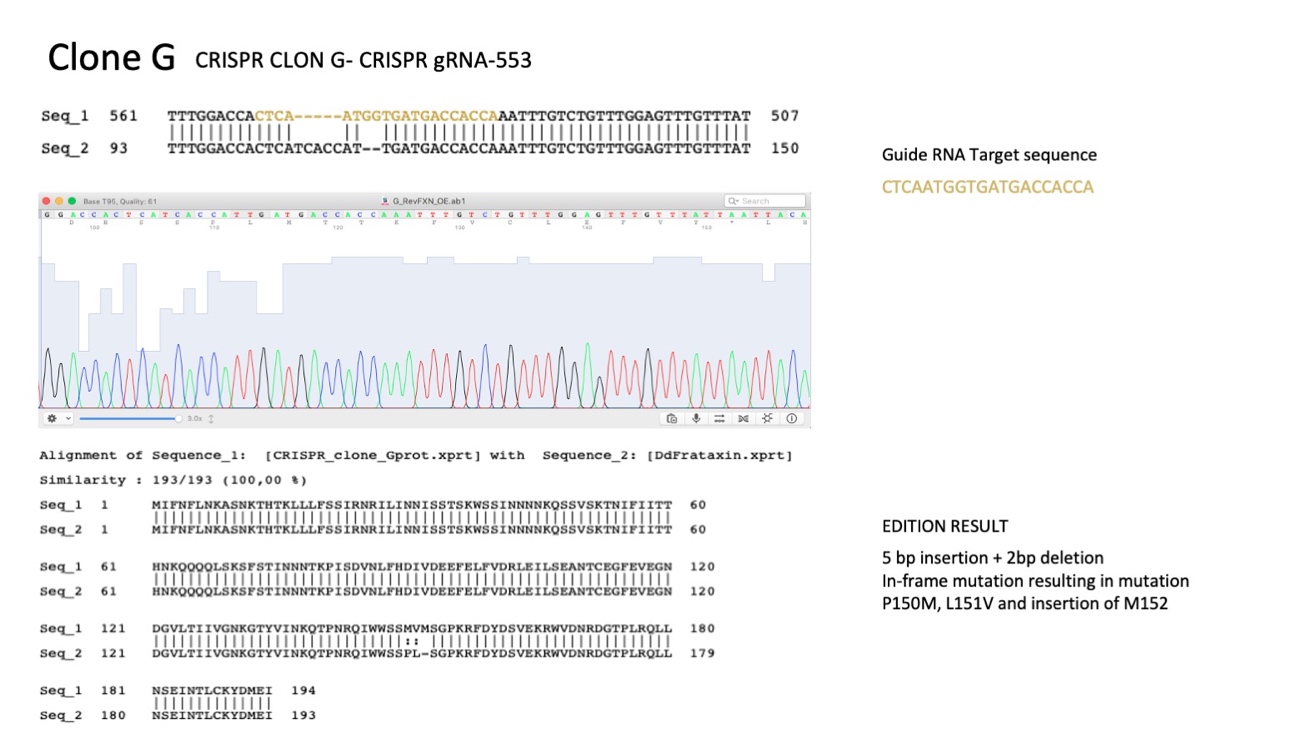

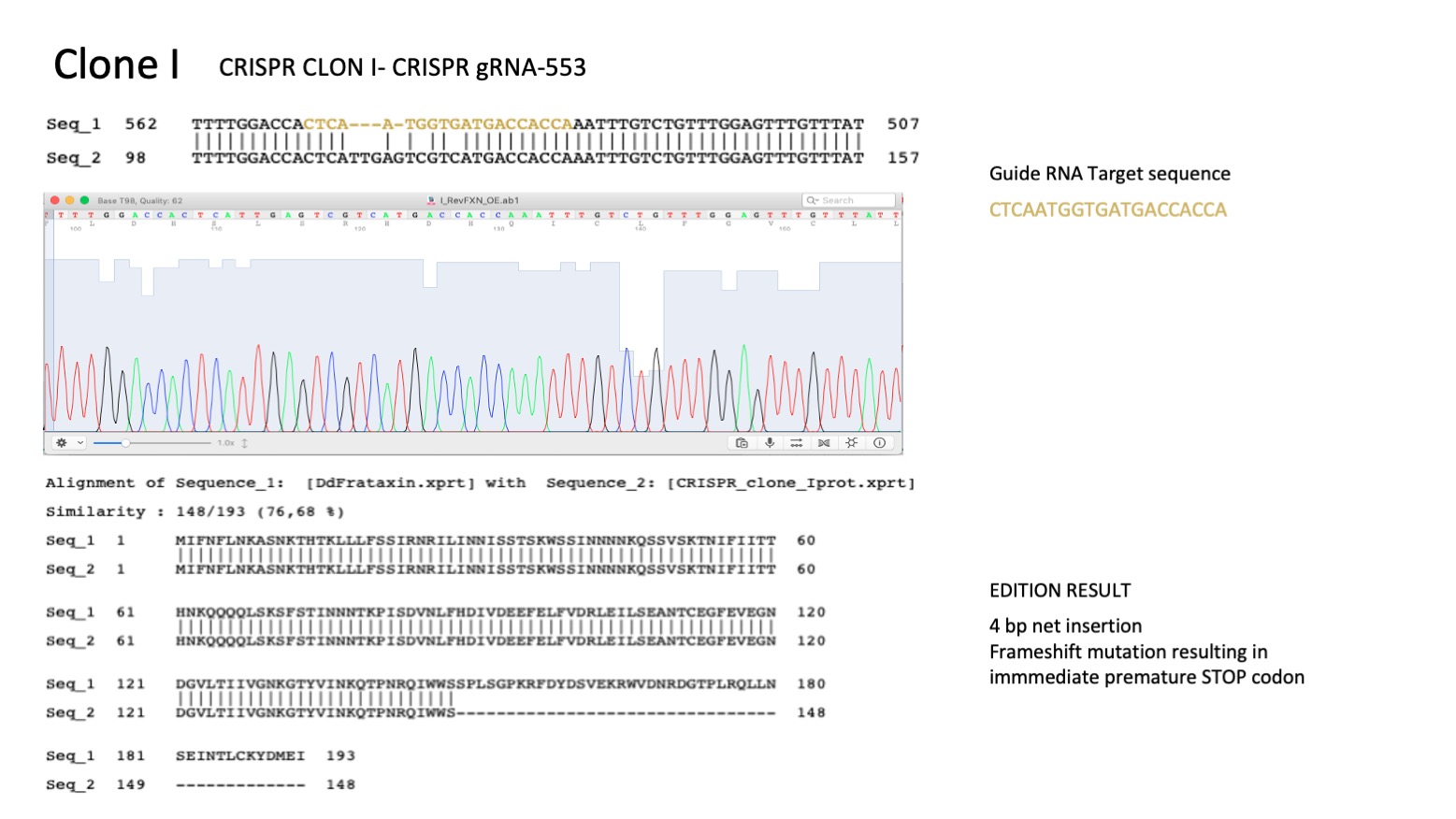

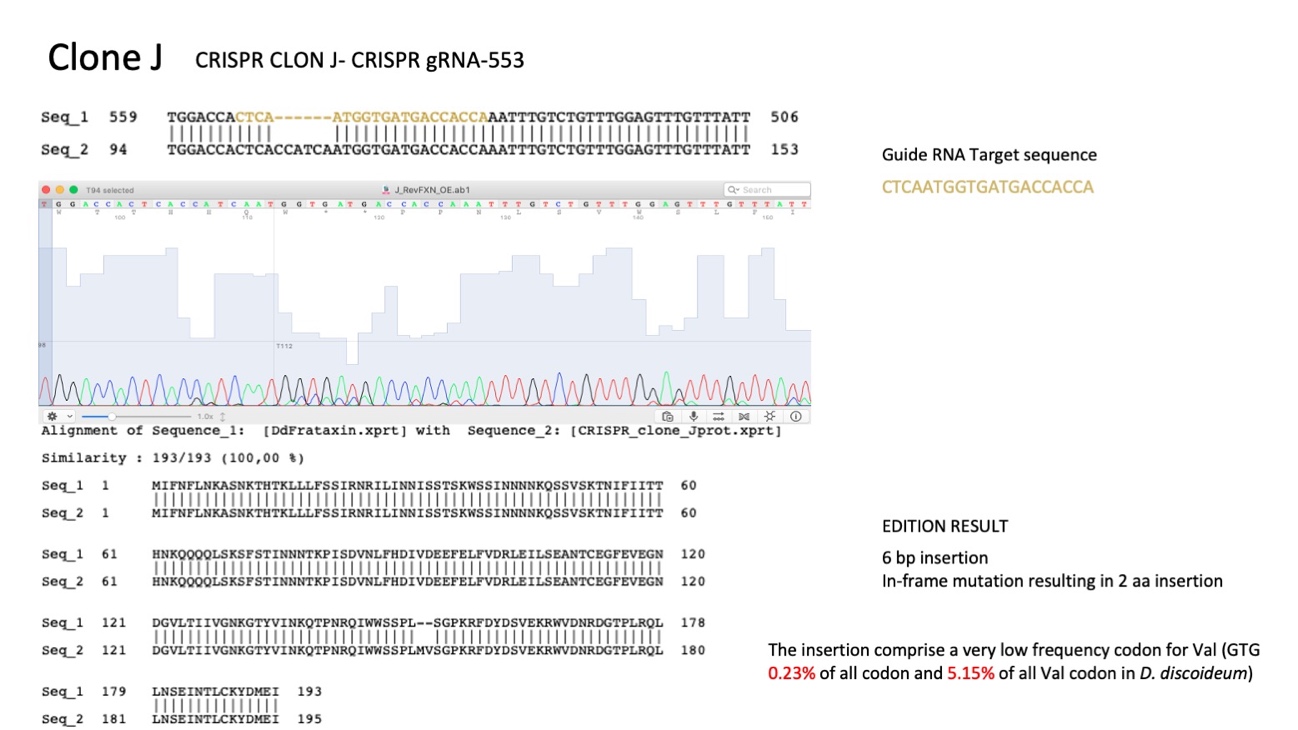

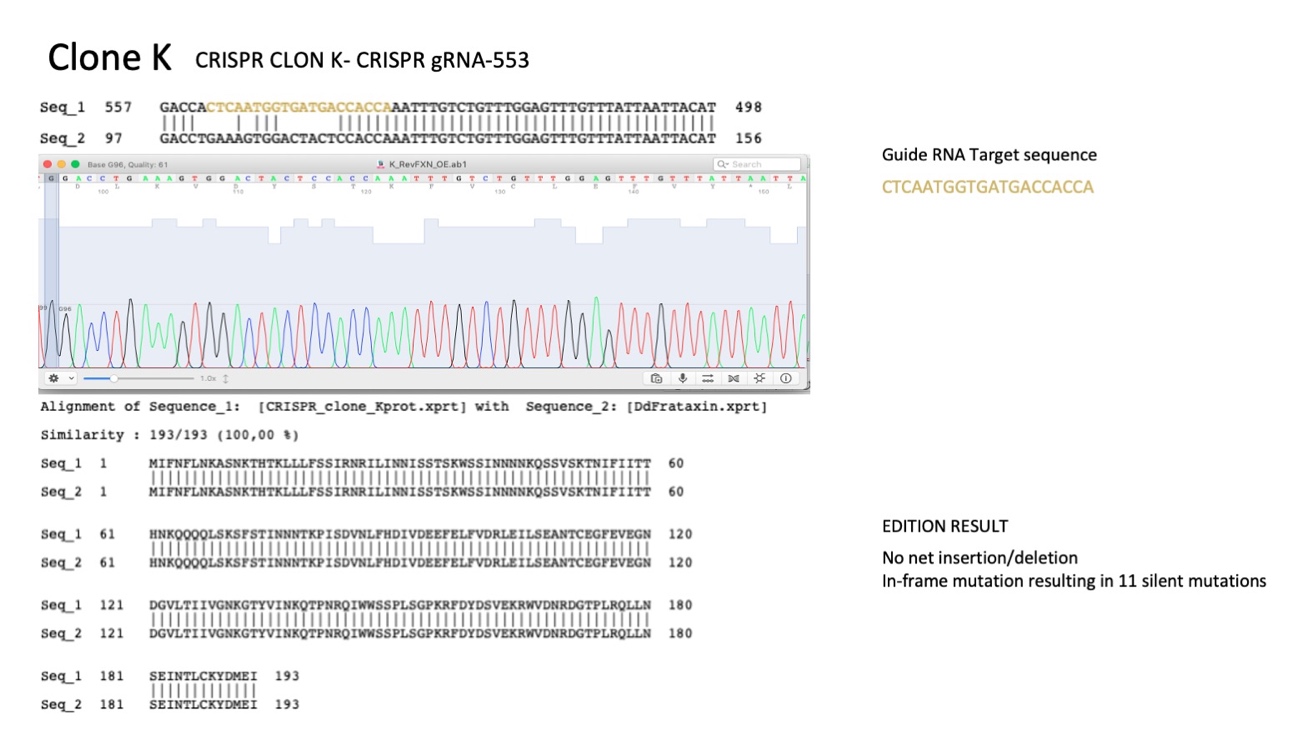

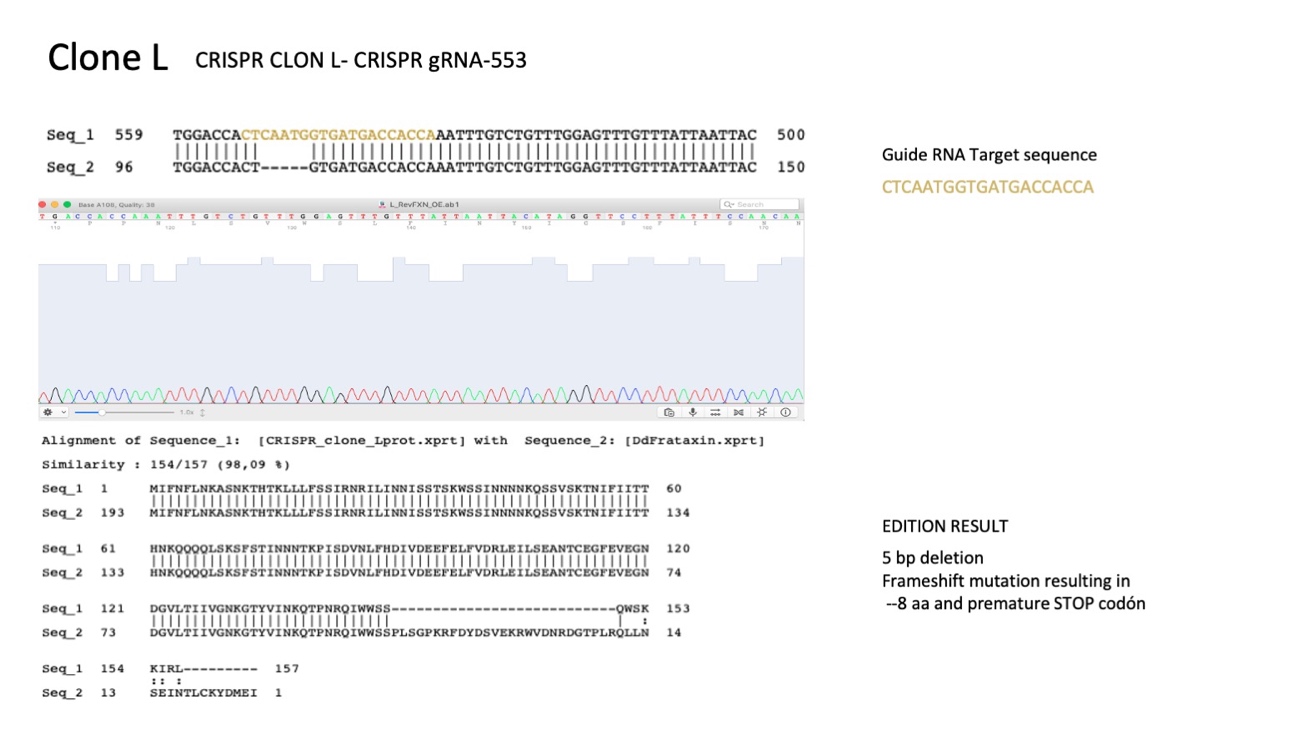

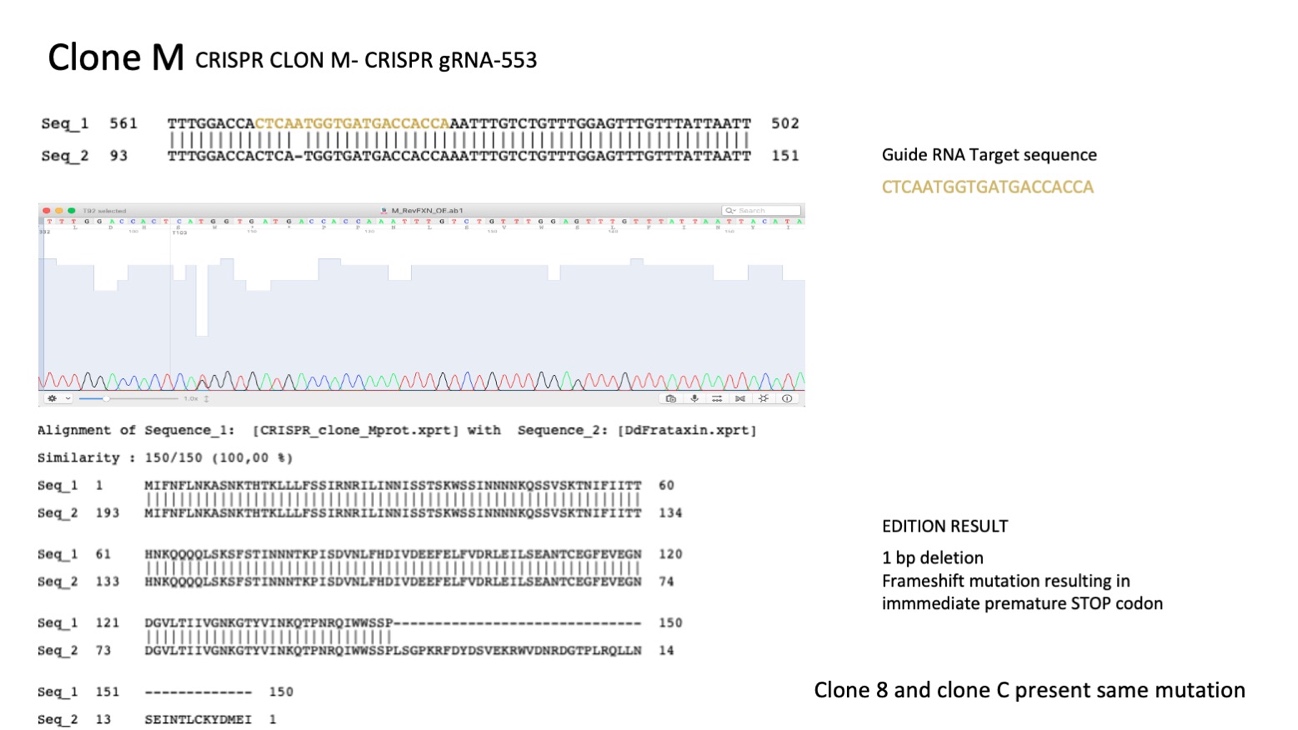

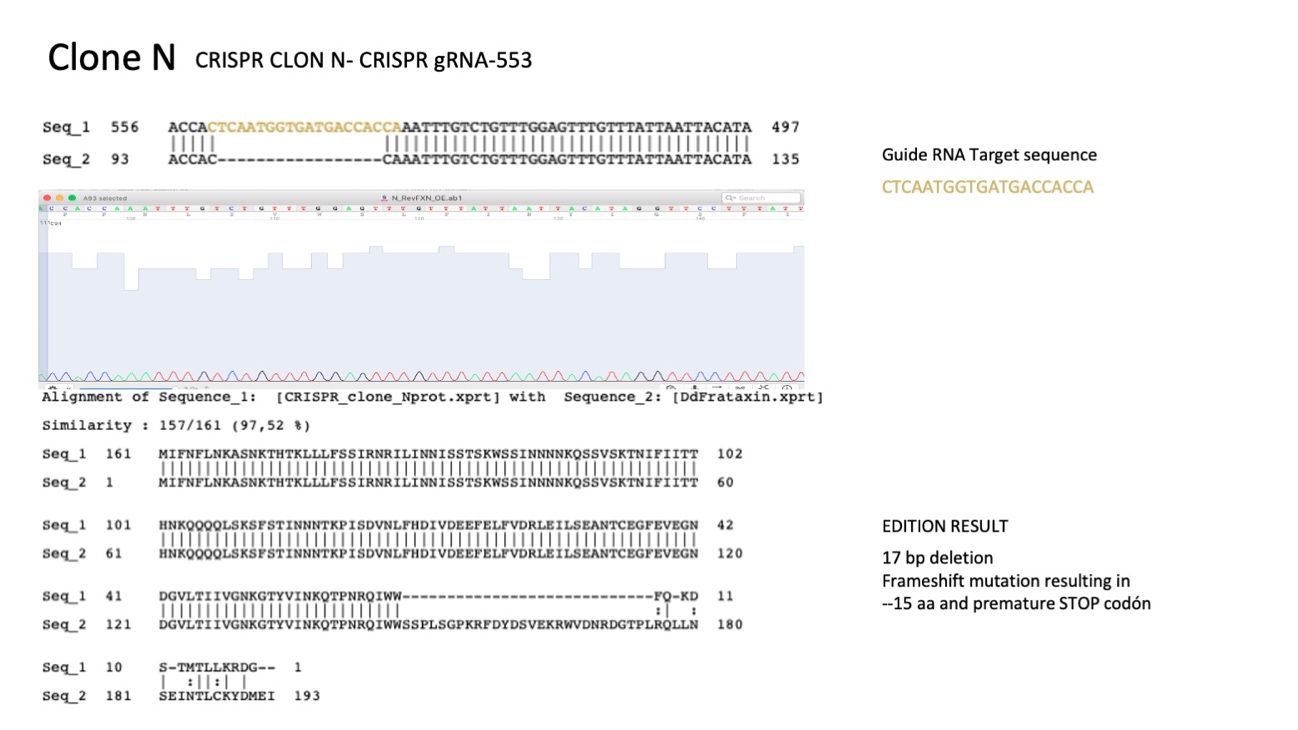

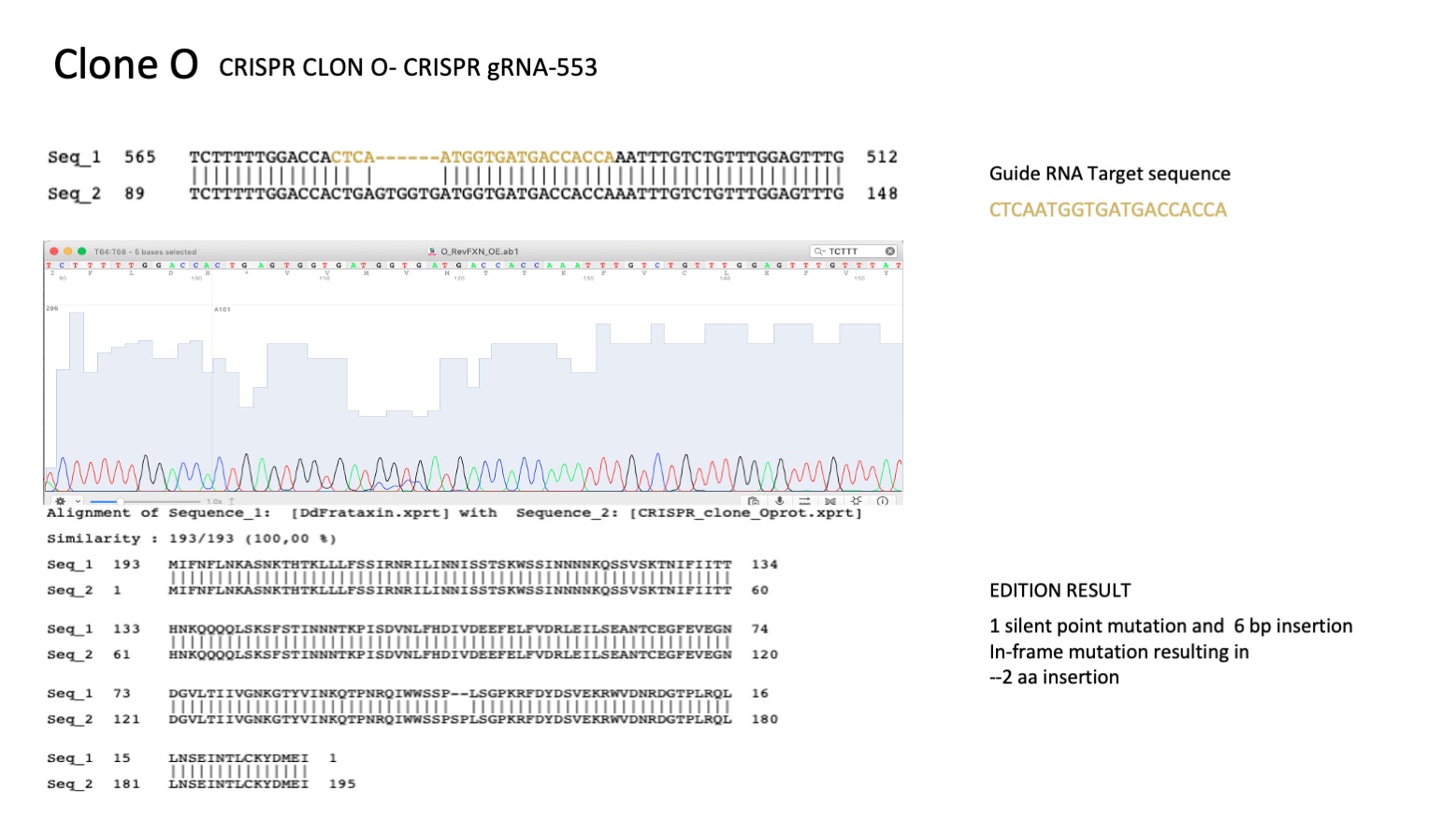

**Figure S2.** Amino Acid Sequence Alignment of *D. discoideum* and human frataxin precursors and predicted fragment from **clone 8**. The black arrow indicates G122 and G130 in the *D. discoideum* and human precursors.

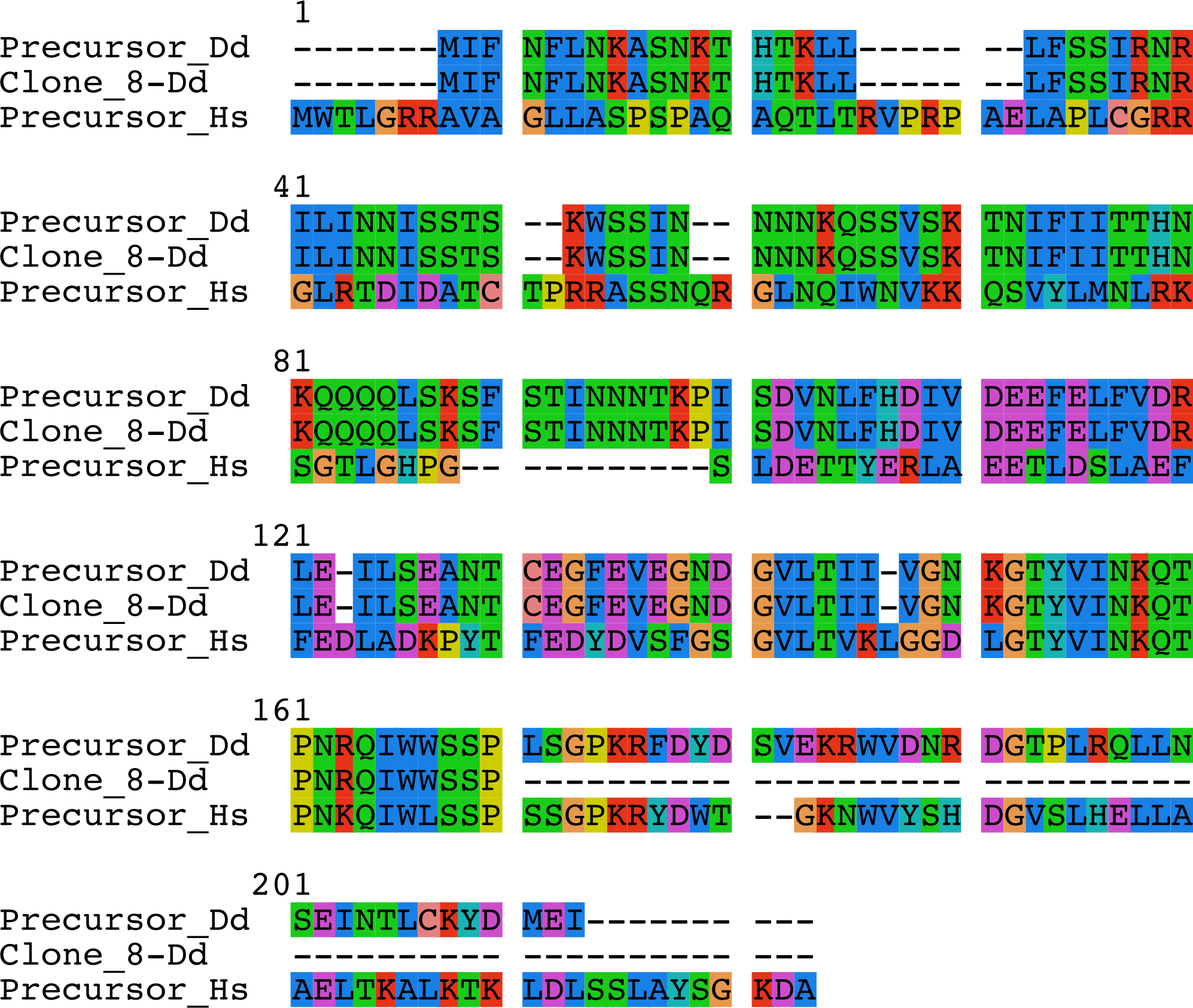

**Figure S3.** Hypothetical dimeric conformation of the truncated form of frataxin found in **clone 8**. Two views are shown. In (A) the predicted intermolecular disulfide bond is shown in sticks and in (B) the N- and C-termini from one subunit are indicated. The model was built by means of AlphaFold2(*52*).

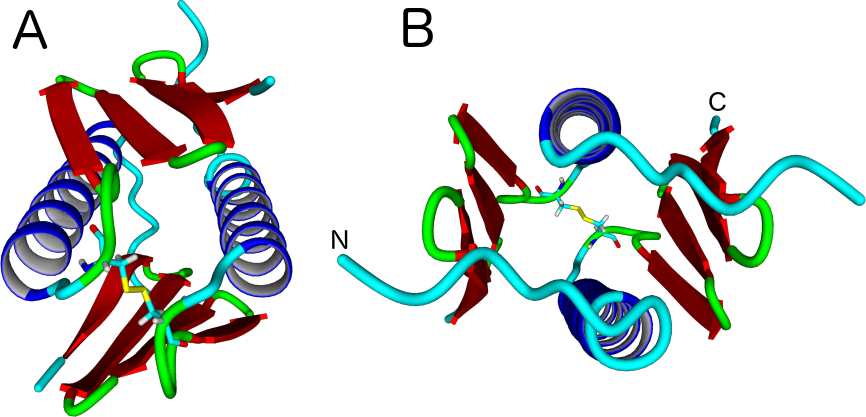

**Figure S4**. **Recombinant Frataxin Detection by Western Blotting Analysis. As** a control experiment, wild-type DdFXN and the fragment expected for clone 8 (residues 81-150) were expressed in *E. coli* and the recombinant protein were detected from the total *E. coli* lysates by NB41 and NB45. Lane 1: Lysate from *E. coli* BL21 (without plasmid). Lane 2: molecular mass markers (also on the right panels, M). Lane 3: lysate corresponding to BL21 cells transformed with pET9A for 81-150 D*. discoideum* frataxin fragment expression (without induction). Lane 4: same as lane 3 but after fragment induction with 1mM IPTG for 3.5h at 37°C. Lane 5: lysate corresponding to BL21 cells transformed with pET9A encoding for the mature 81-193 D*. discoideum* frataxin after induction for 3.5h at 37°C with 1mM IPTG.

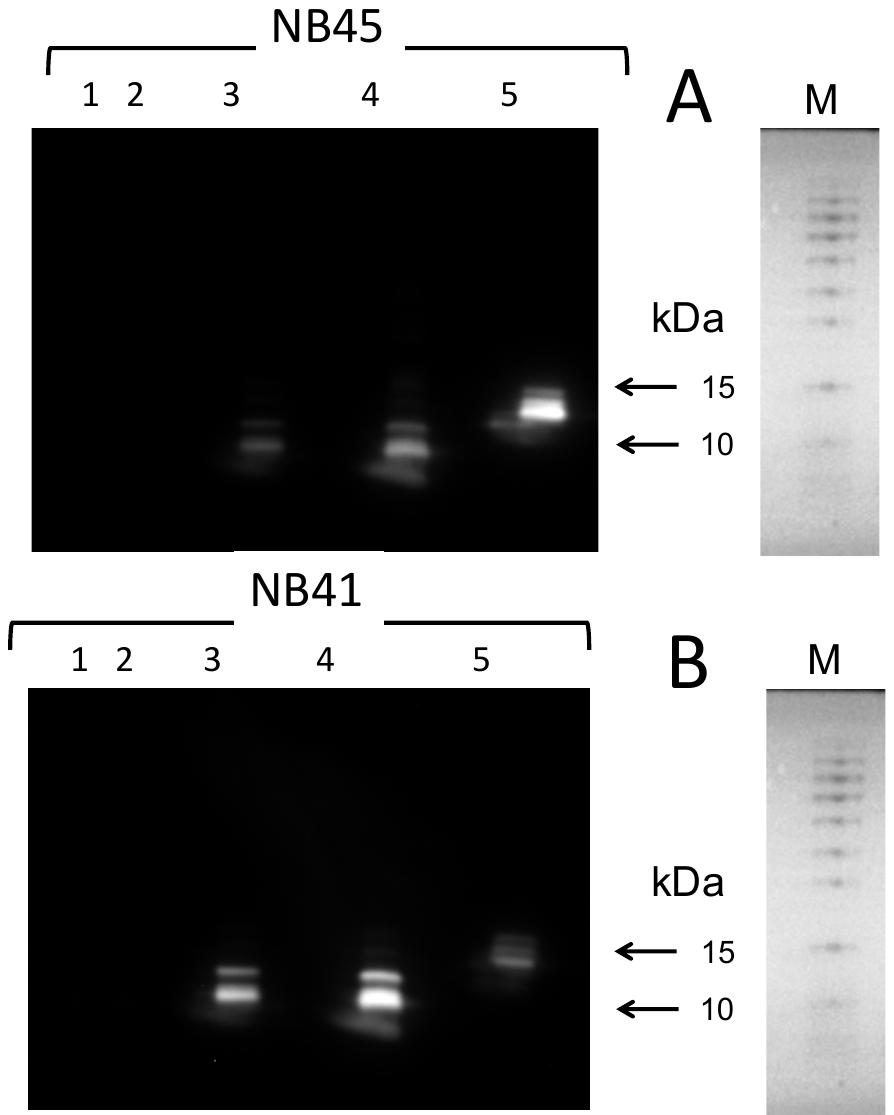
